# Constitutive and inducible fibrosis explain immune variation among threespine stickleback populations

**DOI:** 10.64898/2026.04.27.721125

**Authors:** Emma S. Choi, Ben A. Flanagan, Heather Alexander, John Berini, Alex Yeung, Cole J. Wolf, Viola Watts, Grace Vaziri, Nataly Vargas, Caroline Szajda, Penelope Steffen, Ipsita Srinivas, Mehreen Shahid, Ana Santacruz, Kaligua Rochon, Luke Rippin, Edith Reyes Contreras, Emma S. Redfield, Emma Polard, Cate Patterson, Fahad S. Gilani, Julian Flanagan, Shira Dubin, Peaches Cooper, Panna Codner, Amy Chen, Gwen Casey, Abigail G. Albright, Jessica Hite, Jesse N. Weber, Daniel I. Bolnick, Amanda K. Hund

**Author notes:** Correspondence (E.S.C), (B.A.F.), (D.I.B.), (A.K.H.). Institute for Integrative Cell Biology and Physiology, University of Münster, Münster, Germany (Ana Santacruz). University of Minnesota (Grace Vaziri). These authors contributed equally to this work (Emma S. Choi, Ben A. Flanagan).

## Abstract

Understanding how immune variation arises in natural populations requires disentangling the relative contributions of host genetic differences, environmental variation, and parasite effects, which is rarely possible in wild systems. Threespine stickleback populations vary in their use of intraperitoneal fibrosis as a defense against the helminth parasite *Schistocephalus solidus*, providing a natural system to study the genetic and ecological drivers of immune variation. We combined a 46-lake field survey with common garden experiments on 20 representative populations exposed to multiple parasite genotypes to test whether population differences in fibrosis persist under controlled conditions and whether they depend on parasite genotype or lake ecology. Fibrosis variation was strongly heritable, with both constitutive and inducible components persisting under common garden conditions. In contrast, parasite genotype had only a weak effect on fibrosis responses. Moreover, inducible fibrosis covaried with lake environmental conditions, with populations from more eutrophic-like lakes exhibiting stronger responses than those from more oligotrophic-like lakes. Together, these results reveal ecologically structured divergence in heritable immune responses among natural populations.

## Introduction

Genetic variation in immunity provides the raw material for host defense evolution and contributes to variation in parasite susceptibility among individuals and populations (1–4). Because parasite-driven selection can be strong, immune traits are often highly polymorphic and rapidly evolving (5,6). However, immune variation observed in the wild does not necessarily reflect host adaptive evolution, and may instead arise from phenotypic plasticity, parasite genetic variation, synergistic effects of host and parasite genotypes, or interactions between genotype and the environment. As a result, it remains difficult to determine the extent to which immune differences among populations are genetically based and shaped by natural selection.

Common-garden experiments combined with controlled immune challenges provide a powerful approach for partitioning genetic and environmental contributions to immune variation by minimizing environmental differences and revealing heritable differences among hosts. When paired with data from wild populations, this approach can link immune traits to ecological conditions and potential selective pressures. However, relatively few studies integrate broad sampling of wild populations with experimentally derived estimates of heritable immune variation, limiting our understanding of how immune traits evolve across environmental gradients (7,8). Here, we apply this framework to threespine stickleback (*Gasterosteus aculeatus*), focusing on fibrosis-based resistance to the helminth parasite *Schistocephalus solidus*.

Threespine stickleback and the tapeworm *S. solidus* form a well-studied host–parasite system in which freshwater populations vary in their resistance to infection. Following colonization of freshwater environments by marine ancestors, stickleback populations evolved increased resistance to this parasite (9,10), but now differ markedly in immune responses across lakes (11–14). One key defense is peritoneal fibrosis, a conserved immune response that can encapsulate and restrict tapeworm growth (11,15). Fibrosis reduces parasite success but also imposes fitness costs (16,17), leading to variation among populations in both the presence and magnitude of this response. As a result, some populations exhibit strong fibrosis while others show reduced or absent responses, differences that are observed in both wild and laboratory settings (11,12,14). However, it remains unclear to what extent this natural variation in fibrosis is evolved versus plastic, and whether it is an adaptive response to any environmental variables.

Population-level variation in fibrosis could arise from multiple sources. First, differences may reflect environmental plasticity if populations experience different exposure rates to *S. solidus*, such that fibrosis is induced more frequently or strongly in high-exposure environments; in this case, population differences should disappear under common-garden conditions. Second, variation could be driven by parasite genetic differences, as distinct tapeworm genotypes present different antigens (18) and can elicit variable immune responses in stickleback (19–21), potentially leading to differences in downstream fibrosis induction depending on the tapeworm genotype encountered. Finally, differences may reflect host genetic divergence, with populations evolving differences in constitutive or inducible fibrosis. Constitutive differences should persist in laboratory-reared fish regardless of immune challenge, whereas inducible variation should appear as differences in response to parasite exposure (G×E), potentially modified by host–parasite genotype combinations (G×G).

If fibrosis is heritable, an important question is whether variation is adaptive. Because fibrosis imposes both costs and benefits, its optimal level may differ among populations depending on local conditions (22,23). Stickleback diets vary across lakes: some consume primarily mid-water, planktonic prey (“limnetic”), while others feed mainly on bottom-associated prey (“benthic”) (24–26). Because *S. solidus* is transmitted via planktonic copepods, limnetic-feeding populations are likely exposed to higher tapeworm loads, potentially selecting for stronger fibrosis, whereas benthic-feeding populations may experience weaker selection (27). Alternatively, population differences could reflect non-adaptive processes, such as historical contingency, founder effects, bottlenecks, or mutation accumulation. By linking population-level fibrosis to ecological factors, we can test whether heritable variation corresponds with selective pressures arising from lake-specific environments.

To determine the genetic and ecological drivers of population-level immune variation, we conducted a 46-lake survey of fibrosis in wild stickleback populations. From this, we selected 20 populations spanning the observed range of fibrosis and reared them in a common garden, exposing fish to multiple *S. solidus* genotypes, including local and non-local tapeworms. This design allowed us to test whether population differences persist under uniform conditions, whether they depend on tapeworm genotype, and whether laboratory responses covary with lake ecology. We found that fibrosis variation is heritable, with both constitutive and inducible components, whereas tapeworm genotype had only a weak effect. Inducible responses in the laboratory were correlated with lake ecology, though in an unexpected direction. Overall, this study combines broad population sampling with controlled experimental exposures to disentangle the roles of host evolution, parasite variation, and ecological context in shaping immune variation.

## Materials and Methods

### Field survey of fibrosis and infection

To determine the severity and frequency of the fibrosis phenotype and tapeworm infection prevalence in wild stickleback populations, we sampled lake stickleback populations from three geographic regions on Vancouver Island, British Columbia, Canada (Fig 1); 1) the northern region near Port McNeill, 2) the mid-eastern region near Campbell River, and 3) the west coast region near Bamfield, from lakes listed in Table S1. From May 26 - July 3, 2023 we sampled stickleback from 46 lakes on Vancouver Island using a combination of seining and unbaited minnow traps. For each lake, we quantified mean fibrosis severity and tapeworm infection prevalence using freshly dissected and formalin-preserved individuals (see Table S1 for exact sample sizes). Fibrosis was scored on a 4-point scale, from 0 (no fibrosis), 1 (some fibrosis, organs do not move freely), 2 (fibrosis, organs adhere together), 3 (organs adhered together and to the peritoneal wall), and 4 (severe fibrosis, difficult to open peritoneal cavity) (12). The collections were approved by the Ministry of Forest, Lands, and Natural Resources Operations and Rural Development (Collections permit no. NA23-787881), Fisheries and Oceans Canada (License no. 139753), and the Huu-ay-aht First Nations (Heritage investigation permit no. 2023-013). The sampling sites are located within the traditional territories of the Kwakwaka’wakw and Nuu-chah-nulth First Nations. All collections and experimental activities were approved by University of Connecticut IACUC protocol A21-025.

**Fig 1:**
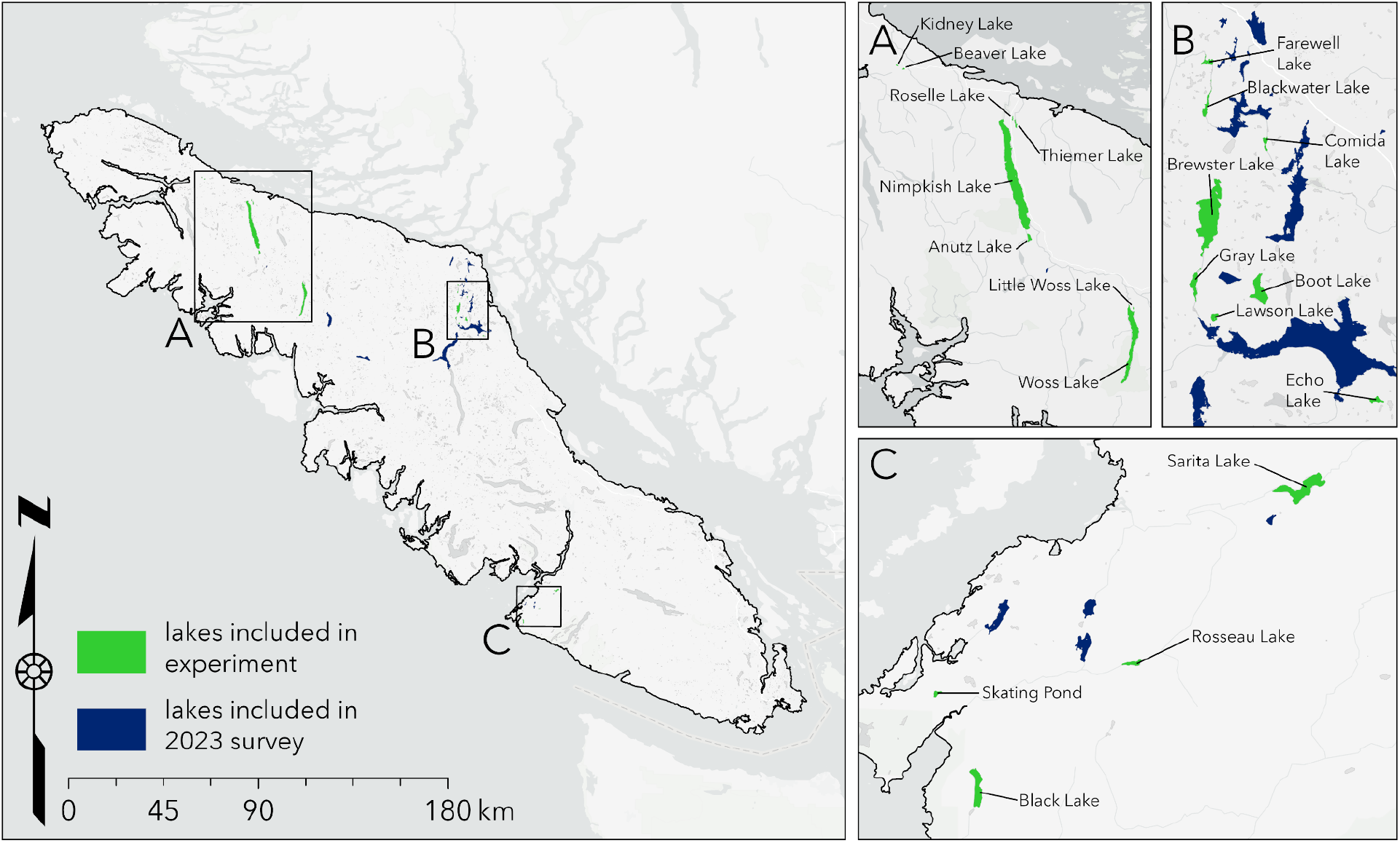
Map of lakes included in the 2023 survey and in the experiment. Lakes included in the experiment were also included in the 2023 survey.

To obtain measurements of lake characteristics, we used an EXO2 multi-probe sonde (YSI Incorporated, Ohio, USA) to measure temperature-depth profiles at one-meter increments to approximately one meter above maximum lake depth. The sonde could measure to a maximum depth of 20 m, and profiles were taken at or near the deepest point of each lake when possible. We also collected multiple water quality metrics including pH, dissolved oxygen (mg L-1; also calculated as % saturation), and chlorophyll *a* (a proxy for phytoplankton biomass). Measurements for phytoplankton provide insight into qualitative differences among lakes and are units of relative fluorescent units (RFUs). Surface temperature was quantified from measurements at the lake surface (less than 5 meters deep), whereas other sonde variables were summarized as averages across all sampled depths.

We obtained lake bathymetric data using iMapBC, an interactive mapping platform maintained by the Government of British Columbia. To extract bathymetry data for each of the focal lakes, we applied specific map layers within the platform, including Bathymetric – 7.5M (under “Base Maps”), and Lake Bathymetric Maps and Digital Bathymetric Maps (under “Fish, Wildlife, and Plant Species”). These layers provided standardized measurements for each sampled lake, including perimeter (m), maximum depth (m), mean depth (m), surface area (ha), and elevation (m).

### Common-garden experiments

In order to determine which factors contribute to variation in wild stickleback fibrosis, we raised stickleback from multiple populations in the lab to minimize environmental variation and eliminate prior exposure to tapeworms. We then tested their fibrosis responses to several immune challenges, including exposure to live tapeworms, injections of tapeworm proteins from different geographic origins, alum injections, and saline injections.

#### Common-garden: animal breeding and rearing

To identify stickleback populations which exhibit variable infection prevalence and fibrosis, we plotted fibrosis against infection prevalence (Fig 2C) and used a preliminary version (Fig S9; reduced sample sizes) to select roughly equal numbers of lakes with high fibrosis and low infection, high fibrosis and high infection, low fibrosis and low infection, and low fibrosis and high infection. This resulted in a subset of 26 of the 46 lake populations sampled in 2023 that were subsequently revisited in June 2024. Using unbaited minnow traps and seine netting, we collected gravid females and mature males to obtain gametes for in vitro fertilization to produce 5 -10 full-sib families per population collected under the Ministry of Forest, Lands, and Natural Resources Operations and Rural Development Collections permit no. NA24-89569, Fisheries and Oceans Canada License no. 139753, and the Huu-ay-aht First Nations Heritage investigation permit no. 2024-042. Gametes for crosses were obtained following the Stickleback IVF Breeding protocol (28). The fertilized eggs were transported to the Bamfield Marine Sciences Center (BMSC), to be reared in aquaria (BMSC AUP #RS-22-09).

**Fig 2:**
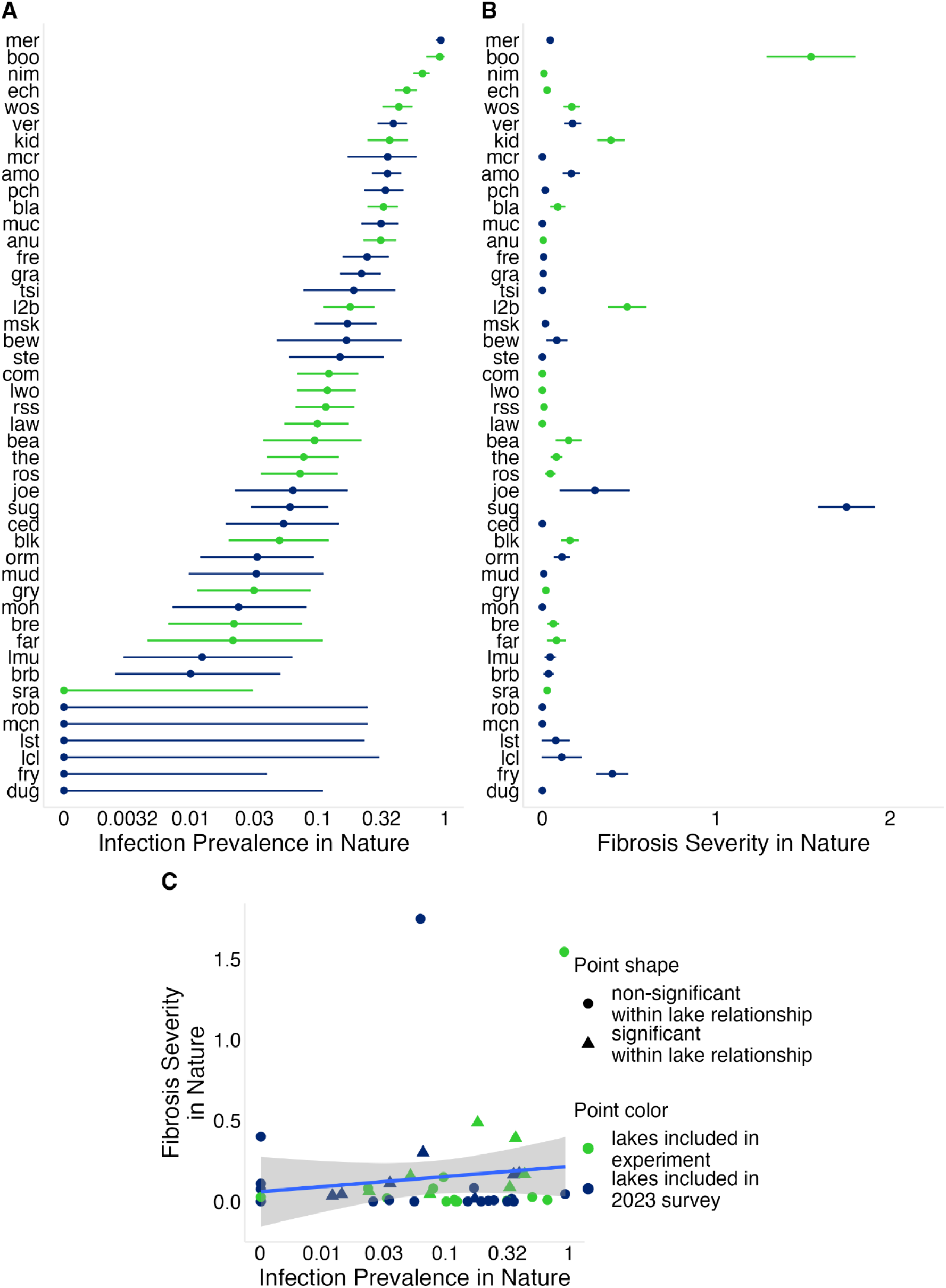
Population-level variation in infection prevalence and fibrosis severity in nature. (A) Infection prevalence across populations;points represent the proportion of infected fish in each population, with 95% binomial confidence intervals. (B) Mean fibrosis severity across populations; points represent population means and error bars show standard errors. (C) Relationship between infection prevalence and fibrosis severity across populations; points represent population means. Triangles indicate populations in which infection and fibrosis are significantly correlated among individual fish (P < 0.05). Infection prevalence in (A) and (C) is plotted on a log scale, with axis labels shown on the original (unlogged) scale. Lake abbreviations correspond to names and sample sizes in Supplementary Table 1.

Once stickleback larvae had consumed their yolk sac, fry were fed live *Artemia* nauplii. When juveniles reached >1.5 cm in length, they were transferred from 3 L Z-hab tanks to 38 L aquaria supplied with sponge filters and artificial plants for enrichment. Fish were fed twice daily with live *Artemia*, and once they reached ∼2 cm in length their diet was supplemented with a ground mixed invertebrate and commercial feed following Stickleback Feeding Protocols (29), along with suspended decapsulated *Artemia* eggs.

We housed the stickleback in an aquarium system with automated flow-through, supplied with fresh water from the Bamfield Water System and dechlorinated using a Waterite™ Excelflow 2472 system (hereafter referred to as “system water”). The light regime was maintained at 16:8 h (light:dark). Because we could not actively control water temperature, it varied seasonally from 15.0 °C to 7.5 °C. Initially, we kept families in the same tank but as the fish grew in size, we occasionally split families among tanks to prevent overcrowding. Fish were maintained under these conditions throughout the experiment.

Once the lab-raised common garden fish were grown, each family was split between treatments to evaluate their fibrosis response, some individuals being exposed to live tapeworms, others injected with antigens to test responses to different immune stimulants, and others were kept as controls.

#### Common garden A: live-tapeworm exposure

To test for population-level differences in infection-induced fibrosis, constitutive fibrosis, and infection rates in lab-raised stickleback, we exposed fish from each population to live tapeworms and quantified fibrosis in exposed fish and unexposed controls. To test for effects of parasite genotype, we used two tapeworm types: Echo Lake tapeworms and Echo Lake × Skogseidvatnet Lake F2 hybrid tapeworms. Tapeworm eggs were generated via a lab-based crossing method described in Weber (9). After collection, eggs were stored in water at 4℃ for at least 4 months. To induce hatching, we distributed eggs across 24-well plates with ∼1mL of water in each well, incubated plates in dark incubators at 18℃ for 1 week, and then exposed plates to 18:6 light:dark cycles at room temperature. Once hatching began (generally within 1-2 weeks after light exposure), we isolated batches of the cyclopoid copepod *Acanthocycops robustus* (∼50 individuals per batch) in ceramic bowls and fasted them for 24 hours. *S. solidus* exposures were performed by pipetting coracidia into the copepod bowls. Although we did not precisely count the number of coracidia, we aimed for >2 tapeworms per copepod (i.e., >100 coracidia per bowl). After 1 week we sampled ∼5 exposed copepods per bowl, anesthetized them in carbonated water, and scanned individuals under a dissection microscope to confirm infection presence. The remainder of the experiment was carried out using copepod cups that were at least 2-weeks post exposure and contained at least one confirmed infection.

We exposed lab-raised stickleback (∼6 months old) to one of three treatments: unexposed controls, native Echo Lake tapeworms (Vancouver Island), or F2 hybrid tapeworms from Echo × Skogseidvatnet Lake (sample sizes in Table S1). Each fish received four copepods drawn from the exposure cups (individual copepods were not screened for infection prior to feeding), and we recorded the number of copepods consumed by each fish; most fish consumed 3–4 of 4 copepods, with no differences among treatments. We maintained the fish for 80 days after exposure before euthanizing them. At that time, we measured each fish’s mass and length (from the caudal peduncle excluding the tail fin to the tip of the lip), scored fibrosis using the 0-4 scale described above, and for exposed fish, recorded the number of encapsulated and free-living tapeworms.

#### Common garden B - Injections

Next, we tested for population-level differences in fibrosis response to an immune stimulant, tapeworm protein, and tapeworm genotypes. Fish from each population were injected with one of four inoculants: 1) endotoxin-free phosphate-buffered saline (PBS) as an injection control; 2) 1% alum solution (10µL of 2% Alumax Phosphate OZ Biosciences and 10µL of endotoxin-free PBS); 3) 0.5 mg/mL tapeworm protein extract. Within each population, half of the individuals receiving tapeworm protein were exposed to local tapeworms from their native region (e.g., northeast, mid-east, or southwest Vancouver Island), and half received foreign tapeworm protein from a different region. Sample sizes are in Table S1.

To obtain tapeworms, we sampled fish from three regions: Anutz (northeast), Boot (mid-east), and Frederick (southwest). Fish were euthanized in MS-222 and frozen at −20°C until dissection. Tapeworms were extracted from the body cavity and homogenized in 0.9× endotoxin-free PBS using a bead beater and 3.2 mm chrome steel beads, then centrifuged at 4000 rpm for 20 min at 4°C to produce protein extract. Protein concentration was measured using a RED 660 kit (G Biosciences) and a PerkinElmer Victor™ plate reader. Extracts were diluted to 0.5 mg/mL in 0.9× PBS and stored at −80°C. Injection solutions were prepared by pooling equal quantities of protein from 4–5 tapeworms from the same lake. Syringes were prepared in a sterile laminar flow hood up to one day before injection and stored at 4°C.

Prior to injection, fish were lightly anesthetized in 50 mg/L MS-222. Pelvic or dorsal spines were clipped to identify treatments; markings were not reused across concurrent experiments. Fish were injected with 20 µL of inoculant into the left peritoneal cavity following Hund et al. (12). The needle was inserted at a shallow angle, and correct injection was confirmed by peritoneal distension. Fish were kept on a damp sponge with gills covered during the procedure (<1 min) and transferred to aerated recovery tanks for ≥10 min. No mortality occurred. Fish assigned to alum and scheduled for day-2 dissection were housed in 38 L tanks; all others were returned to original tanks. Fish from the same family received all treatments and were housed together.

Fish were euthanized 2 or 7 days post-injection. PBS and tapeworm protein groups were dissected at day 7; alum fish were split between day 2 and day 7. During dissection, we measured mass, length, and fibrosis on a 0–4 scale.

##### Quantification and statistical analysis

###### Do infection rates and fibrosis vary among stickleback populations in nature?

We quantified population-level prevalence of *S. solidus* as the proportion of infected individuals per lake. To test whether infection prevalence varied among populations, we used a binomial generalized linear mixed model (GLMM) with lake as a random effect, using 3,321 wild-caught stickleback from 46 populations.

Fibrosis was scored on an ordinal scale from 0 (none) to 4 (severe). We summarized population-level variation by calculating mean fibrosis scores for each lake and tested for covariance with infection prevalence using Pearson correlation.

Because fibrosis scores are ordinal, we analyzed among-population variation using a Bayesian ordinal logistic regression model, following McElreath (30). This approach models each increase in fibrosis score above the preceding fibrosis value, as a set of logistic curves, and estimates the log-odds-ratio (lor) that these increase rates differ among populations, or as a function of some specified treatment. For this analysis, fibrosis scores were shifted into ranks 1-5 to fit software requirements. The Bayesian Ordinal Logistic Regression (BayesOLR) involved modeling an individual’s fibrosis score as an ordered logit function with two sets of parameters: a vector of baseline probabilities of the ranks (‘cutpoints’) and a log-odds-ratio (lor, represented with the symbol ψ) score representing the magnitude of upward transitions through the ranks (e.g., an effect size). That is:

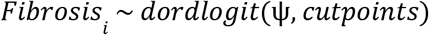

The effect size ψ is then modeled as a linear function of independent variables whose effects we wish to test. For example, to test for among-lake variation in fibrosis for wild-caught fish, we test a model in which:

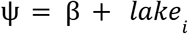

Where β represents the same-wide fibrosis log-odds-ratio (lor), and *lake_i_* is a deviation above or below this mean for any given lake *i*. The overall mean fibrosis is modeled with a normal prior probability distribution with mean = 0 and standard deviation 2 (e.g., β∼*norm*(µ = 0, σ = 2)). The distribution of *lake_i_* effects is modeled as a normal distribution with a mean of zero and a standard deviation that is an estimated parameter: *lake_i_* ∼*norm*(µ = 0, σ = σ*_pop_*). The prior probability distribution for σ*_pop_* is an exponential distribution (with 0 as a lower bound) with mean of 1 (σ*_pop_*∼ *exp*(1)). Among-population variation in fibrosis was assessed using the estimated standard deviation (σ*_pop_*) of lake effects, with larger values indicating greater divergence among populations. Models were fit using the *rethinking* (31) package in R Studio version 2024.12.1+563 (32), with MCMC sampling via the *ulam* function. Posterior distributions were summarized using the *precis()* and *extract.samples()* functions in the *rethinking* package. The cutpoints are modeled with a prior probability distribution that is normal with mean zero and standard deviation of 1.5. For comparison, we also fit a binomial GLMM based on fibrosis presence/absence, which yielded qualitatively similar results but loses resolution due to thresholding; we therefore focus on the Bayesian ordinal model.

###### Is there among-population variation in baseline fibrosis in lab-raised stickleback?

We used a BayesOLR to estimate the magnitude of among-population variation in fibrosis in fish that were not subjected to any immune challenge, not injected with PBS, nor exposed to any tapeworm. This model is the same as the one described above for wild caught fish, with the same prior probability distributions. The specific model used is:

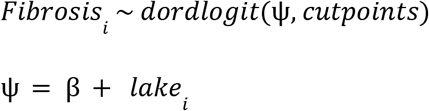

We used a Pearson correlation to test whether Bayesian estimates of baseline fibrosis were associated with Bayesian estimates of fibrosis in natural populations. Lake was treated as the unit of replication, and analyses were conducted on population-level estimates. Likewise, we used a Pearson correlation to test whether Bayesian estimates of baseline fibrosis were associated with Bayesian estimates of fibrosis in saline-injected fish (i.e., whether baseline fibrosis was consistent across control treatments).

###### Is there among-population variation in fibrosis response to alum in lab-raised stickleback?

We used a BayesOLR to evaluate a statistical model where fibrosis depended on population (a random effect), a general response to alum injection (fixed effect), and a population by injection interaction (a random effect). Specifically, we used MCMC (as described above) to search through parameter space to estimate terms in the following model:

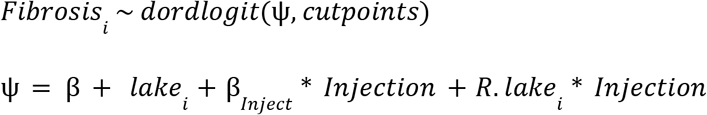

Where Injection is a binary indicator (0 for PBS injected control fish, 1 for alum-injected fish), and *R*. *lake*_i_ estimates the random effect of among-population variation in response to injection (e.g., the population * injection interaction term). For this model we specified the following distributions:

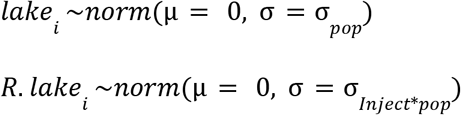

and the following priors:

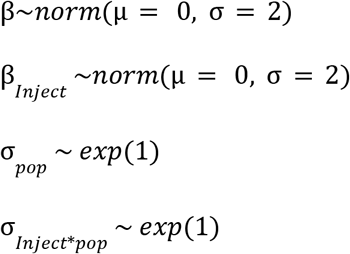

The effect size estimate β*_Inject_* provides a measure of the magnitude and direction of the stickleback response to alum injection (relative to PBS injected controls), regardless of source population. The random effect estimate σ*_Inject_*_\**pop*_ provides a measure of the magnitude of among-population variation in the response to alum injection. Here we are interested in estimating both the overall response effect size and direction β*_Inject_*, and whether σ*_Inject_*_\**pop*_ clearly exceeds zero. We fitted the model separately for fish sampled at 2 and 7 days post-injection, and additionally using a combined model including sampling time as a covariate. To assess consistency of population responses across time points, we correlated posterior estimates of population-specific effects between 2 and 7 days.

###### Do lab-raised stickleback populations exhibit different fibrosis responses to tapeworm protein?

We next analyzed a subset of lab-raised fish injected with PBS (a control), local tapeworm protein, or foreign tapeworm protein. We first tested whether fibrosis increases in response to tapeworm protein (regardless of source) relative to PBS, and whether this response varies among populations. To answer this question we used a BayesOLR with the same model structure as described above for Alum (e.g., ψ = β + *lake_i_* + β*_Inject_*\* *Injection* + *R*. *lake_i_* * *Injection*), but now using *Injection* as an indicator variable contrasting control fish (*Injection* = 0) versus fish injected with either tapeworm protein (*Injection* = 1). We sought to estimate the overall response effect size and direction β*_Inject_*. If the posterior probability distribution for σ*_Inject_*_\**pop*_ clearly exceeds zero, we would infer that lab-raised populations differ from each other in their response to tapeworm antigens.

To test whether responses differ between local and foreign antigens, we fit a second model with *Injection* contrasting foreign (*Injection* = 0) versus local (*Injection* = 1) tapeworm protein. Here, a positive β*_Inject_* indicates a stronger response to local than foreign tapeworms (or vice versa for negative estimates). As above σ*Inject*\**pop* captures among-population variation in this response, with values above zero indicating population differences in response to local-foreign antigen responses (e.g., a genotype by genotype interaction effect).

###### Do lab stickleback populations exhibit different fibrosis responses to tapeworm exposure?

We analyzed fibrosis responses to controlled exposures of live tapeworms in lab-raised fish using three related Bayesian ordinal logistic regression models. First, we separately analyzed responses to Echo Lake tapeworms and F2 hybrid tapeworms using the same model structure as above:

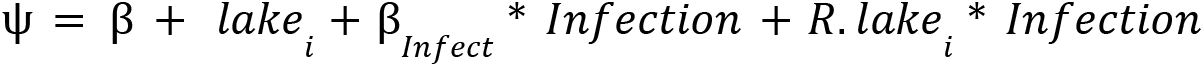

Here *Infection* is a binary indicator (0 for unexposed control fish, 1 for fish that were fed a controlled dose of tapeworm infected copepods). As before we wish to estimate the magnitude and direction of the effect of exposure (β*_Infect_*), and whether this effect varies among stickleback populations (σ*_Infect_*_\**pop*_>> 0). We analyzed Echo and hybrid tapeworm exposures separately because they differed in infection dose (number of infected copepods per fish). We then tested whether population-specific responses were correlated between the two datasets. Finally, we fit a third model comparing exposure to Echo versus hybrid tapeworms directly, to test whether responses differed by tapeworm source and whether this varied among populations. For this third analysis the *Infection* term is a binary indicator (0 for Echo Lake exposures, 1 for hybrid exposures).

###### Do laboratory measures of fibrosis response covary with lake ecological variables?

To determine if constitutive or inducible fibrosis is correlated with environmental covariates, we first conducted a principal component analysis using average lake temperature less than 5 meters deep, average lake pH, average lake chlorophyll *a*, log10 max lake depth, and log10 lake area (Table S3 shows loadings). We then used a Pearson correlation to determine if environmental PC1 was correlated with either constitutive or inducible fibrosis.

Sample sizes for all statistical tests are provided in Table S1 and other statistical parameters are available in the results section. All statistical analyses were performed in the R statistical environment (32), using the *ggplot2* (33) package for plotting. All data and R scripts are available on a data repository.

## Results

### Population differences in infection prevalence and fibrosis in nature

We surveyed 46 Vancouver Island lake stickleback populations in 2023 to quantify variation in fibrosis severity and *S. solidus* infection prevalence (Fig 1; Table S1). Infection prevalence varied widely among lakes, ranging from 0% to 93% across populations (Fig 2A). A binomial GLMM estimated mean prevalence at 10.5%, with strong among-population variation (ICC = 0.49), indicating nearly half the variation in infection was attributable to lake identity.

Fibrosis severity also differed among populations (Fig 2B). We found strong among-population variation (σ*_pop_* = 1.93 [1.41, 2.65]), with 22% of variance attributable to lake differences. A linear model gave similar support (F₄₅,₁₅₃₂ = 19.12, P < 0.0001). Sugsaw (sug) and Boot (boo) lakes showed the highest mean fibrosis (1.75, se = 0.16; 1.55, se = 0.25, respectively), whereas 12 lakes showed no visible fibrosis.

Lakes with higher log *S. solidus* prevalence tended to have higher fibrosis, but the relationship was weak and non-significant across lakes (r = 0.14, P = 0.37; Fig 2C), consistent with previous results from Alaska (14). By contrast, within lakes, infected individuals had higher fibrosis than uninfected fish (lco = 1.82 [1.19, 2.35]), consistent with previous laboratory and field studies showing infection-associated induction of fibrosis (11). The magnitude of this effect varied among populations (σ*_pop_* = 0.71 [0.18, 1.39]), indicating that lake identity modulates the strength of the infection–fibrosis relationship. However, this variation in infection-associated fibrosis explains only a fraction of the among-lake variation in fibrosis severity.

### Common garden experiment: heritable variation in fibrosis

#### Populations differ in constitutive fibrosis

We observed population differences in baseline (constitutive) fibrosis in lab-reared stickleback that did not receive immune stimulation (Fig 3A). In the uninjected controls, fibrosis was generally low (mean of 0.44), but showed substantial among-population variation (σ*_pop_*= 1.89 [1.15 , 2.91]; Supplementary Data 2, Panel D), indicating heritable differences in constitutive fibrosis. Five populations showed no fibrosis, while Boot Lake (boo) showed the highest mean fibrosis (2.30, se = 0.169). Fibrosis in PBS-injected controls was highly correlated with uninjected fish across populations (r = 0.76, P = 0.0001), indicating minimal effect of injection itself. Mean fibrosis in the wild was positively associated with baseline fibrosis in the lab from the same populations (Fig 3B; r = 0.51, P = 0.022), although this relationship weakened after excluding Boot Lake (Fig S2; r = 0.35, P = 0.12).

**Fig 3:**
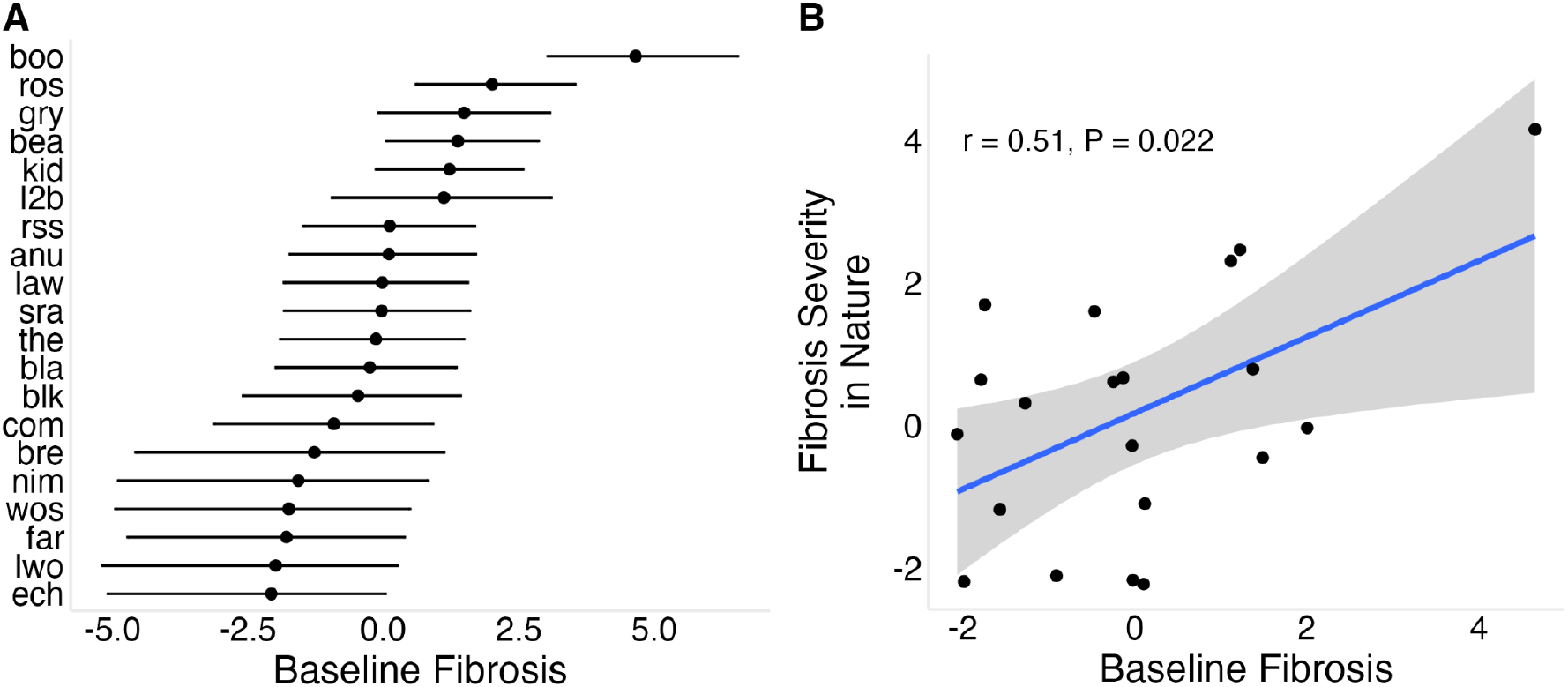
Population-level variation in baseline fibrosis and its relationship to wild fibrosis. (A) Baseline fibrosis across populations; points represent Bayesian estimates for uninjected control fish, standardized such that 0 represents the mean across populations. (B) Relationship between Bayesian estimates of baseline fibrosis measured in the laboratory and fibrosis severity observed in wild populations.

#### Populations differ in their fibrosis response to an adjuvant (alum)

To determine if populations vary in the rate of fibrosis induction following immune stimulation, we exposed fish to alum and measured responses at 2 and 7 days post-injection (dpi). At 2 dpi, alum injection increased fibrosis relative to saline controls (lco = 1.07 [0.60 , 1.53], Supplementary Data 2, Panel E), but there was no detectable variation among populations in this response (σ*_pop_* = 0.3 [0.02 , 0.84], Supplementary Data 2, Panel F). By 7 dpi, the effect of alum injection was stronger (lco = 2.46 [1.88 , 3.02], Supplementary Data 2, Panel G), and among-population variation in response increased as well (σ*_pop_* = 0.45 [0.02, 1.18], Supplementary Data 2, Panel H). Using both timepoints in a combined model, we estimated the per-day increase in the odds of being in a higher fibrosis category following alum injection, which revealed both a strong overall effect of alum injection (lco = 0.42 [0.33 , 0.50], Supplementary Data 2, Panel I) and significant among-population variation in the rate of response (σ*_pop_* = 0.11 [0.03 , 0.22], Supplementary Data 2, Panel J). This indicates that populations differ in the speed of fibrosis induction following immune stimulation. Representative populations showed distinct temporal trajectories of fibrosis induction (Fig 4A).

**Fig 4:**
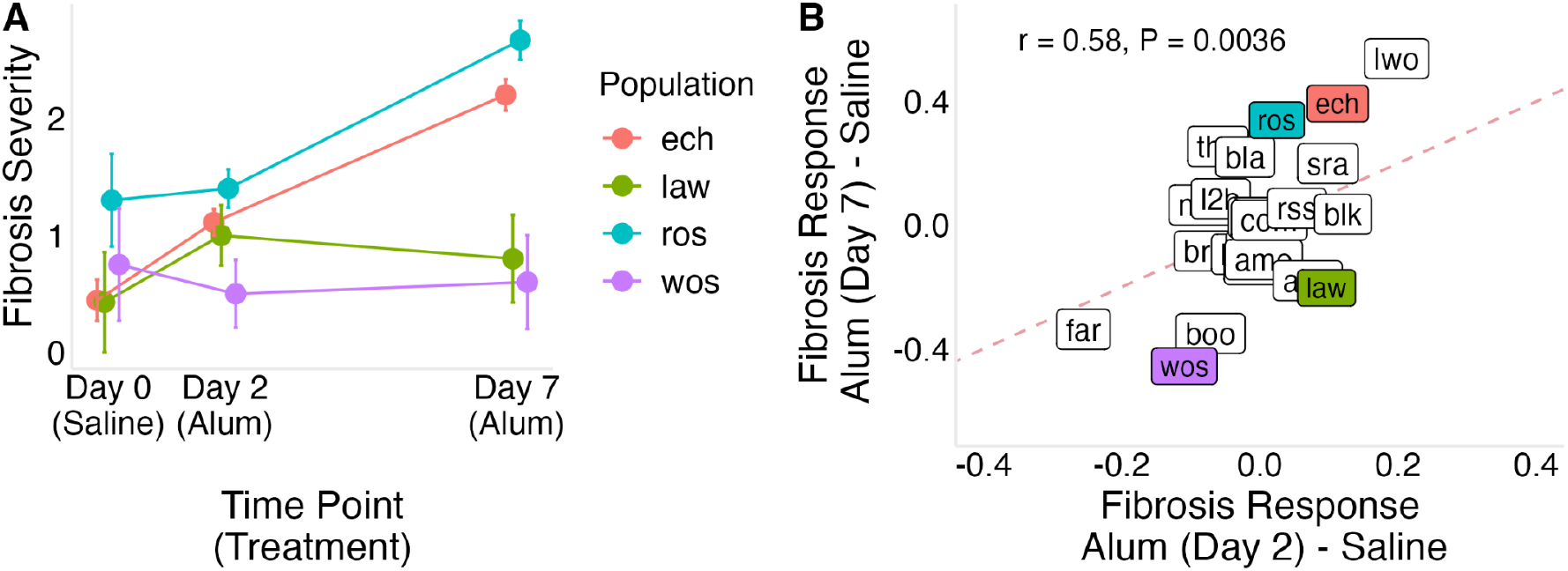
Temporal dynamics of fibrosis responses to alum injection across populations. (A) Fibrosis severity in saline-injected controls (day 0), and alum-injected fish at two and seven days post-injection, shown for four representative populations. (B) Fibrosis response Bayesian effect sizes (alum-injected compared to saline control) at two and seven days post-injection; points represent population-level responses. Colored points correspond to the populations highlighted in (A): Roselle Lake (ros) shows a stronger increase from day 2 to day 7, Echo Lake (ech) shows similar responses at both time points, Lawson Lake (law) shows a rapid early response with little subsequent increase, and Woss Lake (wos) shows minimal change. The dashed line indicates equal responses at day 2 and day 7.

#### Tapeworm genotype affects fibrosis during live infection but not protein injection

To test whether tapeworm genotype influences host fibrosis responses, we injected lab-reared fish with proteins derived from geographically local and distant *S. solidus* populations. Injection with tapeworm protein produced a weak overall effect on fibrosis (lco = 0.33 [-0.11 , 0.75], Supplementary Data 2, Panel K), but there was moderate evidence for among-population variation in response (σ*_pop_* = 0.47 [0.06 , 1.05], Supplementary Data 2, Panel L), indicating population differences in sensitivity to tapeworm antigens. There was no overall effect of tapeworm genotype on fibrosis (lco = 0.07 [-0.37 , 0.48], Supplementary Data 2, Panel M), and only moderate support for among-population variation in differential responses to local and foreign tapeworm proteins (i.e. a G×G effect; σ*_pop_* = 0.33 [0.03 , 0.89], Supplementary Data 2, Panel N). Nevertheless, responses to locally and geographically distant tapeworm proteins were strongly correlated across populations (r = 0.58, P = 0.007, Fig 5A), suggesting limited divergence in antigen-specific induction.

**Fig 5:**
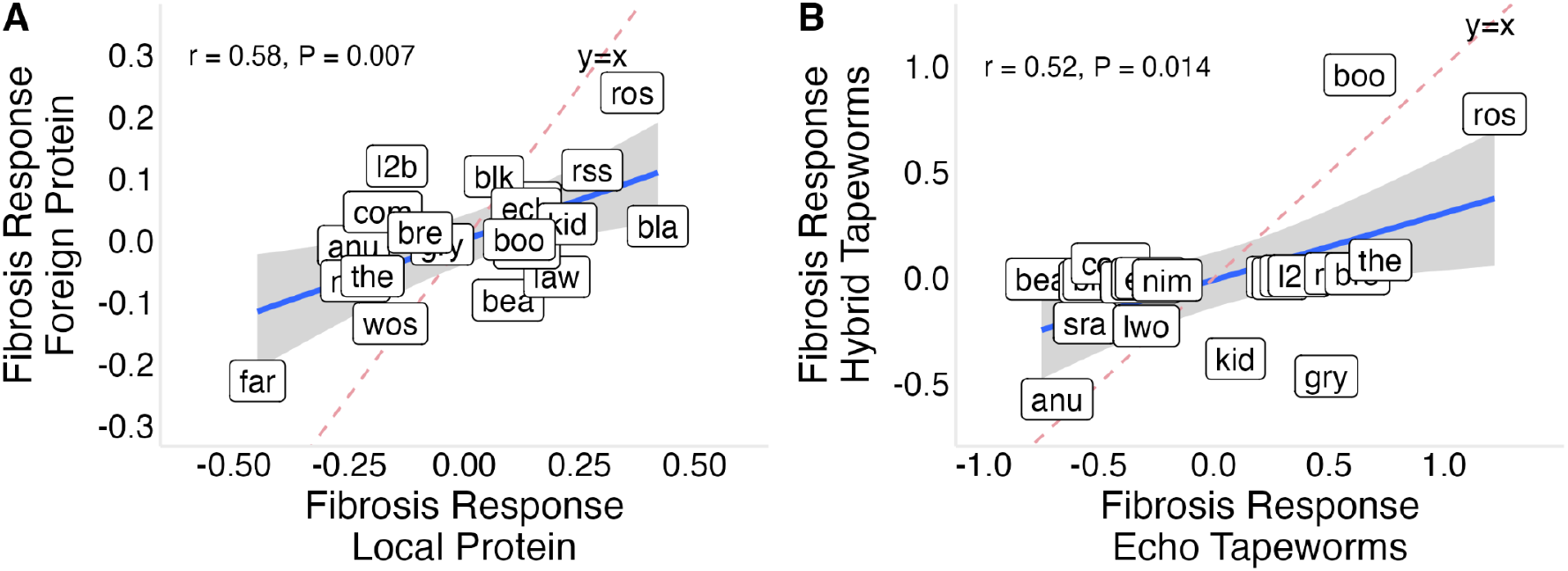
Population-level fibrosis responses to local versus foreign tapeworm exposures. (A) Fibrosis responses to local versus foreign tapeworm protein injection across populations; points represent population-level Bayesian posterior estimates. Responses are positively correlated, indicating no consistent effect of tapeworm genotype in protein exposure. (B) Fibrosis responses to live Echo versus hybrid tapeworm exposure across populations; points represent population-level Bayesian posterior estimates. Responses are not correlated across populations, and Bayesian ordinal logistic regression supports an effect of tapeworm genotype during live infection.

To test whether tapeworm genotype effects extend across broader geographic scales, we experimentally exposed lab-reared stickleback to live *S. solidus* from a local Vancouver Island population (Echo Lake) or to hybrid tapeworms combining local and geographically distant lineages (Echo Lake × Skogseidvatnet Lake, Norway). Lab-raised fish exposed to native Echo Lake tapeworms showed lower fibrosis than fish exposed to hybrid tapeworms (Echo - hybrid lco = -0.85 [-1.89 , -0.07], Supplementary Data 2, Panel O). Although the magnitude of this genotype effect varied among stickleback populations (σ*_pop_* = 1.09 [0.25 , 2.22], Supplementary Data 2, Panel P), responses to Echo and hybrid tapeworm exposure were positively correlated across populations (r = 0.52, P = 0.014; Fig 5B) indicating that populations differ primarily in the strength, rather than the direction, of their responses to tapeworm genotype.

Interpretation of genotype effects in live infections is limited by differences in exposure intensity between treatments: Fish were exposed to live tapeworms via infected copepods, but the infection rate in copepods differed between treatments, leading to higher effective exposure to Echo Lake tapeworms. This difference would be expected to produce stronger fibrosis in response to Echo tapeworms. However, we instead observed higher fibrosis following exposure to hybrid tapeworms that include foreign genotypes from Europe. This pattern provides further support for an effect of tapeworm genotype on host immune activation, although interpretation remains limited by unequal exposure intensity between treatments. Given these differences, subsequent analyses focus on fish exposed to Echo Lake tapeworms.

#### Heritable variation in inducible fibrosis responses among stickleback populations

To characterize heritable variation in fibrosis response among stickleback populations, independent of tapeworm genotype effects, we used results from the live tapeworm exposures and tapeworm protein injections described above. For live Echo tapeworm exposures, infection success was low across populations (5.6% [3.17%, 9.06%], Fig S7) and showed no consistent association with environmental covariates (Fig S8), consistent with these hosts having low susceptibility to exposure. Stickleback exposed to Echo Lake tapeworms did not show elevated fibrosis relative to unfed controls, with a weak overall trend in the opposite direction (lco = -0.77 [-1.85 , 0.2], Supplementary Data 2, Panel Q). Despite this, there was support for a population by exposure interaction effect (σ*_pop_* = 0.92 [0.06, 2.18], Supplementary Data 2, Panel R), indicating that populations differ in their response to tapeworm exposure, reflected in the distribution of Bayesian estimates of population-specific responses to tapeworm exposure (Fig 6A). A similar pattern was observed for hybrid tapeworm exposure, which showed no overall effect on fibrosis (lco = 0.18 [-0.61, 0.92], Supplementary Data 2, Panel S), but comparable among-population variation in response (σ*_pop_* = 0.91 [0.25, 1.58], Supplementary Data 2, Panel T). Although exposure effects for individual populations were highly uncertain, both assays consistently indicated non-zero among-population variation in responsiveness to infection.

**Fig 6:**
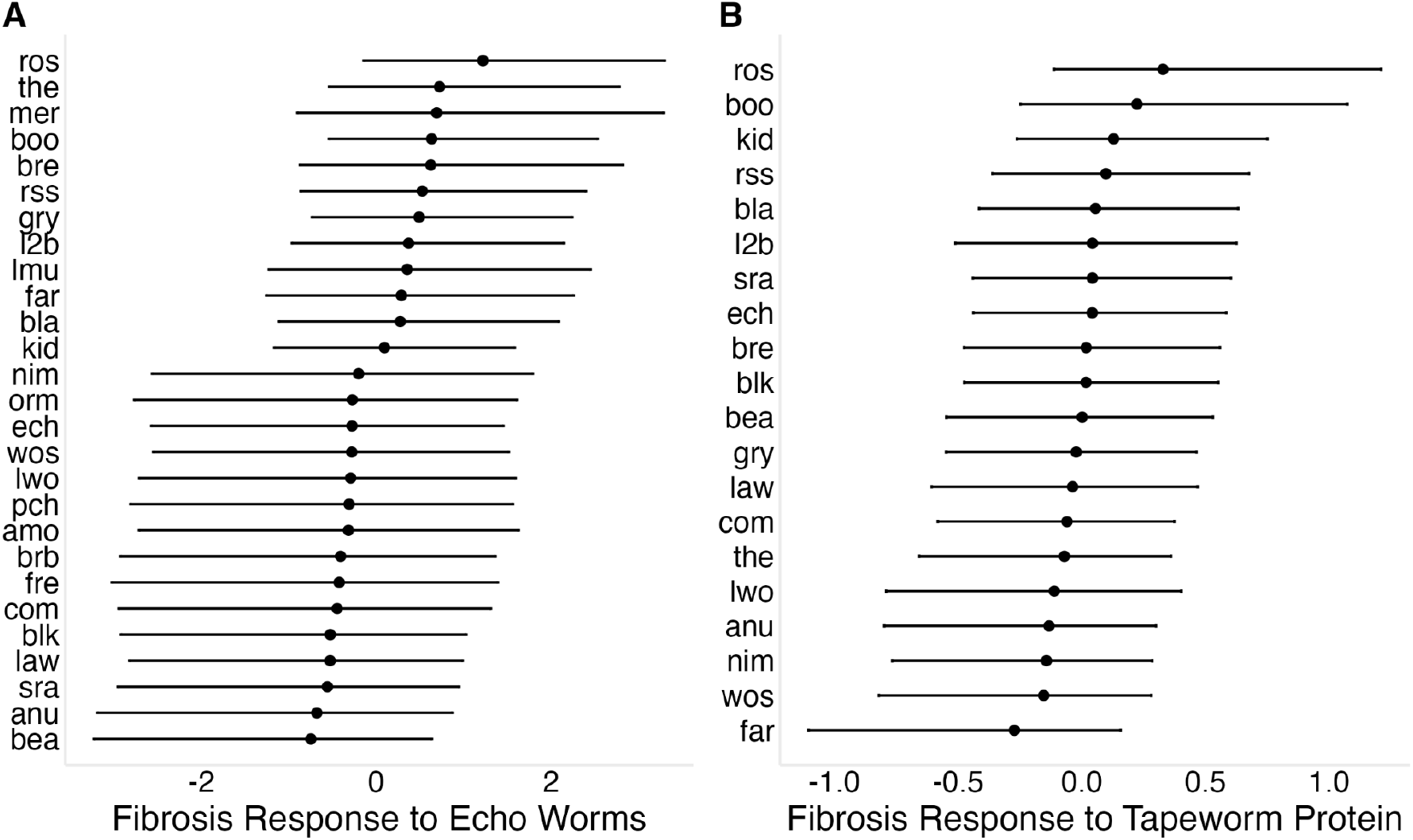
Population-level variation in fibrosis responses to tapeworm exposure. (A) Fibrosis response to live Echo Lake tapeworm exposure; points represent population-level Bayesian posterior means and error bars show 95% credible intervals. (B) Fibrosis response to tapeworm protein injection; points represent population-level Bayesian posterior means and error bars show 95% credible intervals.

Because live tapeworm exposures integrate multiple processes (e.g. infection success, parasite growth, immune activation by the host, and immune suppression by live tapeworms), we used tapeworm protein injections to isolate antigen-driven fibrosis responses. Tapeworm protein produced a modest overall increase in fibrosis (lco = 0.38 [-0.04, 0.78], Supplementary Data 2, Panel K), with evidence for among-population variation in response (σ*_pop_* = 0.25 [0.02 , 0.66], Supplementary Data 2, Panel L), reflected in a broad distribution of Bayesian estimates of population-specific responses to tapeworm protein injection (Fig 6B). Raw mean differences alongside population-level Bayesian estimates (with credible intervals) are provided in Table S2 to aid interpretation of population-specific responses, which ranged from negligible to strong induction, and raw fibrosis scores are displayed in Figure S1. Protein- and live-exposure responses were positively correlated (r = 0.48, P = 0.03; Fig S3C).

To test whether laboratory immune phenotypes predict fibrosis in wild populations, we related Bayesian estimates of population effects for wild fibrosis to corresponding estimates of constitutive and inducible (population × treatment) fibrosis responses measured in the laboratory. Constitutive fibrosis explained 26% of variation in wild fibrosis estimates, compared to 15% for tapeworm protein responses and 11% for alum responses. However, these relationships were not robust to exclusion of Boot Lake (boo); without Boot, explained variance dropped to ∼4–6% across predictors, indicating that associations were strongly influenced by a single high-leverage population.

Population constitutive fibrosis was positively correlated with inducible responses to tapeworm protein (r = 0.70, t = 4.20, P = 0.0006), but not with alum responses at day 2 (r = −0.05, t = −0.21, P = 0.84) or day 7 (r = −0.23, t = −0.99, P = 0.34). By contrast, inducible responses were only weakly and non-significantly correlated across challenges (tapeworm protein vs alum day 2: r = 0.25, t = 1.11, P = 0.28; tapeworm protein vs alum day 7: r = 0.26, t = 1.14, P = 0.27).

We also tested whether laboratory assays can explain among-lake variation in fibrosis-infection associations observed in the wild - that is, the extent to which infected fish show elevated fibrosis relative to uninfected fish within a lake. This association was consistently positive but varied in magnitude among populations. We tested whether this variation could be explained by population-specific laboratory estimates of fibrosis responsiveness to immune stimulation. Among-lake variation in fibrosis–infection associations showed a weak positive relationship with laboratory responses to tapeworm protein (r = 0.290, t = 1.28, P = 0.218), and was not associated with responses to alum or Echo Lake tapeworm exposure (both P > 0.5). Responses to Canada–Norway hybrid exposure showed a marginal positive correlation (r = 0.42, t = 2.09, P = 0.05).

### Inducible fibrosis responses vary with lake ecology

Constitutive fibrosis showed no significant association with environmental PC1 (58.6% of variance explained; Fig S5A; r = 0.16, P = 0.51), but this relationship was sensitive to population composition: excluding Boot Lake (boo) yielded a positive trend (Fig S5; r = 0.45, P = 0.06), indicating dependence on a high-leverage population. In contrast, inducible responses showed a clearer association with environmental variation (Fig 7; r = 0.53, P = 0.02), which was qualitatively consistent but attenuated in Bayesian estimates due to shrinkage (Fig S4; r = 0.35, P = 0.14). Individual covariates showed similar patterns, with lake depth, area, and pH negatively associated with inducible fibrosis (P < 0.05, Fig S6). Together, these results suggest that lake ecology is more consistently associated with inducible than constitutive fibrosis, with stronger inducible responses in more eutrophic systems.

**Fig 7:**
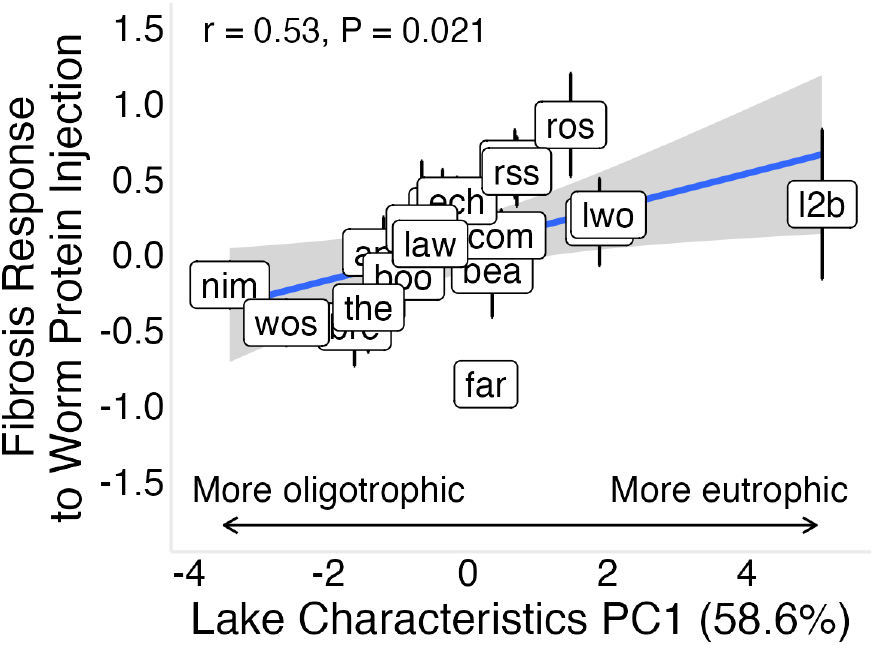
Association between environmental variation and inducible response to tapeworm protein. Fibrosis response to tapeworm protein injection is positively associated with lake environmental PC1 across populations. Response is calculated as the raw difference in fibrosis score between tapeworm protein–injected and control fish. PC1 represents a trophic gradient, separating more oligotrophic lakes (deeper, higher oxygen, lower chlorophyll *a*) from more eutrophic lakes (shallower, lower oxygen, higher chlorophyll *a*). A similar but weaker positive trend is observed using Bayesian population-level estimates (Fig S3).

## Discussion

To determine the sources of immune variation among wild stickleback populations, we used a common garden experiment to distinguish among alternative explanations for population differences in fibrosis, including environmental plasticity, tapeworm genotype effects, and heritable divergence in immune traits. We found the strongest support for heritable differences among populations, with evidence for variation in both constitutive and inducible fibrosis, and limited support for plastic or parasite genotype explanations.

These results extend previous work identifying genetic variation in fibrosis among a small number of populations (11,12)), and show that such variation is more widespread across lakes than previously appreciated. Importantly, we find that both constitutive and inducible components of fibrosis vary among populations, with constitutive fibrosis accounting for more of the observed variation across populations. However, this pattern is influenced in part by a single high-leverage population. In contrast, inducible responses show more consistent associations with ecological conditions than constitutive levels.

To understand how these components of fibrosis relate to one another, we considered covariation among constitutive and inducible responses across immune challenges. Constitutive fibrosis was positively associated with inducible responses to tapeworm protein, and protein- and live infection responses were correlated. In contrast, responses to alum were not consistently correlated with tapeworm-based assays, indicating partial independence between general inflammatory responses and parasite-specific immune pathways.

### Little support for non-genetic explanations for fibrosis variation

If population-level variation in fibrosis were driven solely by differences in tapeworm exposure, we would expect minimal among-population differences under controlled laboratory conditions. Instead, substantial variation in fibrosis persisted among populations reared without immune stimulation as well as those exposed to alum, tapeworm proteins, and live tapeworms. This suggests that exposure-driven plasticity alone does not fully account for observed population differences in the wild.

A second non-host-genetic explanation is that variation reflects differences in tapeworm genotype across populations. Under this scenario, we would expect consistent population-specific responses to local versus foreign tapeworm antigens, and for fibrosis patterns in the field to correspond to responses induced by sympatric parasites. However, this expectation is not supported within Vancouver Island: populations from the same geographic region, likely exposed to similar tapeworm genotypes due to bird-mediated dispersal (34), exhibited marked differences in fibrosis, and tapeworm genotype had no detectable effect on among-population variation. Together, these results suggest that neither exposure history nor parasite genetic variation can fully explain the observed population-level differences at this spatial scale.

At broader geographic scales (North America vs Europe), however, parasite genotype did influence responses: fibrosis was lower following exposure to local Echo parasites than to hybrid tapeworms. This pattern is consistent with local parasites evading or suppressing host immunity, whereas foreign parasites are more readily detected, echoing a “resist globally, infect locally” dynamic (9).

### Constitutive genetic differences in fibrosis among lake populations

Fibrosis differed among stickleback populations raised in a common lab setting, confirming fibrosis variation is highly heritable. We find evidence that this variation occurs both constitutively and in the magnitude of inducible response to immune challenge. Fibrosis varied extensively among populations in control fish (i.e., unexposed or saline-injected), suggesting heritable differences in constitutive fibrosis among populations. These differences were broadly consistent with variation observed in wild-caught fish, although the strength of this correspondence was influenced by a single high-leverage population. These results are notable because stickleback fibrosis has generally been considered an inducible trait, activated in response to immune stimulation (11,12,15,19).

This raises the question of why some populations exhibit constitutive fibrosis despite its known fitness costs. Elevated constitutive fibrosis may be favored under high *S. solidus* pressure if it reduces the costs of repeated inflammatory responses (35), if populations evolve to overshoot optimal defenses (36), or if it provides protection against a broader suite of parasites (37,38). However, constitutive fibrosis observed in the laboratory does not always match patterns in wild populations; some populations exhibited elevated constitutive fibrosis under laboratory conditions despite showing little to no fibrosis in wild-caught individuals. One explanation is that laboratory environments, with reduced pathogen exposure and altered microbial communities, may induce hyper-reactive immune states and elevated baseline fibrosis (39,40).

### Genetic variation in inducible fibrosis responses

Fibrosis also differed between control fish and those challenged with immune stimuli (alum, live tapeworms, or tapeworm protein). The response to alum was especially strong, consistent with the activation of an evolutionarily ancient pathway predating jawed vertebrates (15). Responses to live tapeworms and tapeworm protein were generally weaker, although we still observed among-population variation in laboratory settings, consistent with heritable differences in inducible fibrosis (GxE interactions).

We expected inducible fibrosis to reflect alternative defense strategies, including unresponsiveness in naïve marine populations , inducible resistance (11,12,41), or tolerance via suppression of fibrosis (11). The populations exhibiting weak or absent inducible fibrosis responses could reflect limited evolution of inducible resistance following freshwater colonization or alternative defense strategies that reduce reliance on fibrosis.

In the Nimpkish (nim) and Anutz (anu) Lake populations, stickleback morphologically resemble marine fish, retaining extensive armor plating (Yeung and Bolnick, unpublished). Because armor plate reduction is a rapid and well-characterized marker of freshwater adaptation (42,43), the persistence of marine-like morphology in these populations suggests limited freshwater adaptation likely due to ongoing gene flow with marine populations. These marine-like populations fail to mount fibrotic responses following tapeworm exposure and show limited constitutive fibrosis, consistent with previous inferences that marine stickleback lack fibrosis (11,14).

At the other extreme, low inducible fibrosis in some populations is consistent with tolerance. In Gosling Lake, tolerance is associated with a deletion in the immune regulatory gene *spi1b*, reducing fibrosis potential and pro-fibrotic activity (11,44). However, tolerance may not always be favored: immune-capable immigrant genotypes nearly replaced Gosling Lake ancestry within ∼10 generations, suggesting costs of tolerance even in locally adapted populations (45). These contrasting outcomes indicate that alternative defense strategies are dependent on ecological context which is substantiated by our observation that reaction norms are related to ecological covariates.

### Environmental and ecological drivers of the fibrosis reaction norms

The strong relationship between the inducible fibrosis response and environmental conditions indicate this type of response plays an outsized role in the eco-evolutionary dynamics of tapeworm defenses (Fig S10). Fibrosis responses were strongest in smaller, shallower lakes with higher chlorophyll *a* and lower dissolved oxygen (environmental PC1; Fig 7), providing the first direct evidence linking immune variation to lake ecology.

Stickleback diet varies seasonally and is associated with ecomorph, with limnetic fish generally consuming more copepods and often carrying higher parasite burdens within lakes (27,46–48). However, across lakes, benthic populations exhibit higher average infection loads than limnetic populations, a countergradient pattern attributed to stronger immunity in limnetic fish (27). Consistent with this, stickleback from benthic zones tend to have higher tapeworm abundance than those from limnetic zones (49).

One explanation for our results is that benthic populations may consume fewer copepods overall but encounter more heavily infected copepods, leading to higher effective exposure in smaller, oligotrophic lakes. Alternatively, limnetic populations may experience higher encounter rates but evolve tolerance, although this would predict higher infection prevalence than we observe. More plausibly, inducible fibrosis is favored in oligotrophic lakes where copepods, though a smaller dietary component, are more likely to carry infection.

Supporting this, tapeworm eggs deposited by aquatic birds hatch preferentially in shallow, warm, sunlit waters (50), increasing effective transmission in shallow lakes by expanding the spatial extent of suitable habitat and concentrating hosts, parasites, and intermediate hosts. These processes likely increase encounter rates between stickleback and infected copepods in shallow systems, elevating selection for inducible fibrosis. Although lake ecology could directly influence immunity, our results more strongly support an indirect mechanism mediated by parasite exposure through the copepod community.

## Conclusion

To capture immunological diversity in nature, it is important to sample multiple populations, as laboratory studies and single-population designs can obscure natural variation in immune traits and their ecological and evolutionary drivers (51). By integrating field surveys with controlled laboratory challenges across many populations, we linked genetic and environmental variation to immune phenotypes in wild stickleback.

Wild populations exhibited substantial variation in fibrosis. In common garden conditions, baseline fibrosis explained a major component of among-population differences, highlighting constitutive variation as an underappreciated axis of immune differentiation. Some populations consistently showed little or no fibrosis, consistent with reduced resistance or alternative strategies such as tolerance. In contrast, inducible fibrosis was strongly associated with lake characteristics, indicating that local ecology shapes immune responses. These results show that constitutive immune variation represents a major axis of population differentiation, whereas inducible variation is more tightly linked to environmental conditions, illustrating how host genetics and ecology jointly structure immune traits in the wild.

## Data Availability

Dissection data from the field survey and experiment have been deposited at Zenodo as 10.5281/zenodo.19824354 and are publicly available as of the date of publication. All original code has been deposited at 10.5281/zenodo.19824354 as of the date of publication.

## Author contributions

Conceptualization: DIB, AKH. Resources: DIB, AKH, JH, JW. Data curation: EC, BF, JB, JH, IS. Methodology: AKH, DIB, HA. Funding acquisition: AKH, DIB, JH. Supervision: DIB, AKH. Formal analysis: EC, BF, DIB. Investigation: HA, BF, EC, JB, GV, NV, AS, CS, PS, MS, ER, CP, FG, JF, SD, PC, ERC, AC, GC, AA, DIB. Visualization: BF, EC, JB, DIB. Project administration: HA, VW, EP, KR. Writing - original draft: EC, BF. Writing - review & editing: EC, BF, DIB.

## Supporting information

Supplementary_Data2_Bayesian_Posterior_Outputs

## Acknowledgements

This research was conducted on the traditional territories of the Kwakwaka’wakw and Nuu-chah-nulth First Nations. Thanks to Sara Lane, Jürgen Ehlting, and John Taylor at the University of Victoria for support during protein extractions. The Bamfield Marine Sciences Center provided invaluable staff support and facilities during the course of the experiment. During the preparation of this work the author(s) used ChatGPT to assist with editing and refinement of language and clarity. After using this tool/service, the author(s) reviewed and edited the content as needed and take(s) full responsibility for the content of the published article.

