## Supplementary_Data1_Figures S1-S10_TableS3 for "Constitutive and inducible fibrosis explain immune variation among threespine stickleback populations"

Supplementary Information 1

Contains:

- Supplementary Figures S1–S10
- Table S3

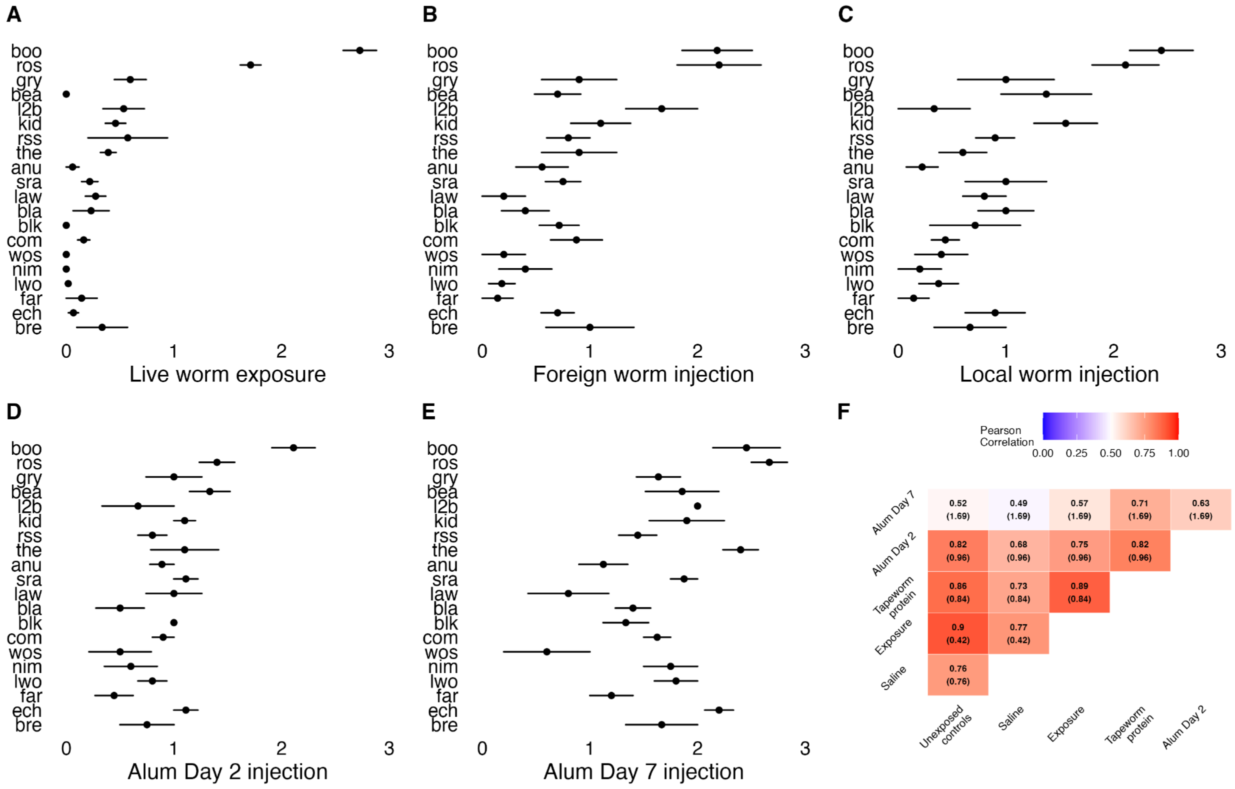


Figure S1: Variation in fibrosis severity for A) fish exposed to live tapeworms, B) fish injected with foreign tapeworm proteins, C) fish injected with local tapeworm proteins, D), fish injected with alum and dissected 2 days post injection, E) fish injected with alum and dissected 7 days post injection, and F) the pairwise correlation coefficient among treatments with correlation coefficient and the mean fibrosis score for the treatment row.

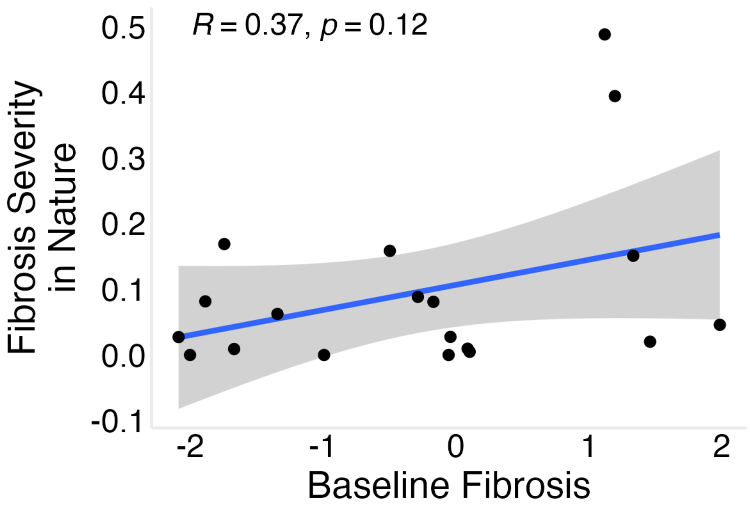


Figure S2: Correlation between Bayesian estimates of baseline fibrosis and fibrosis severity in nature, excluding Boot Lake from the analysis.

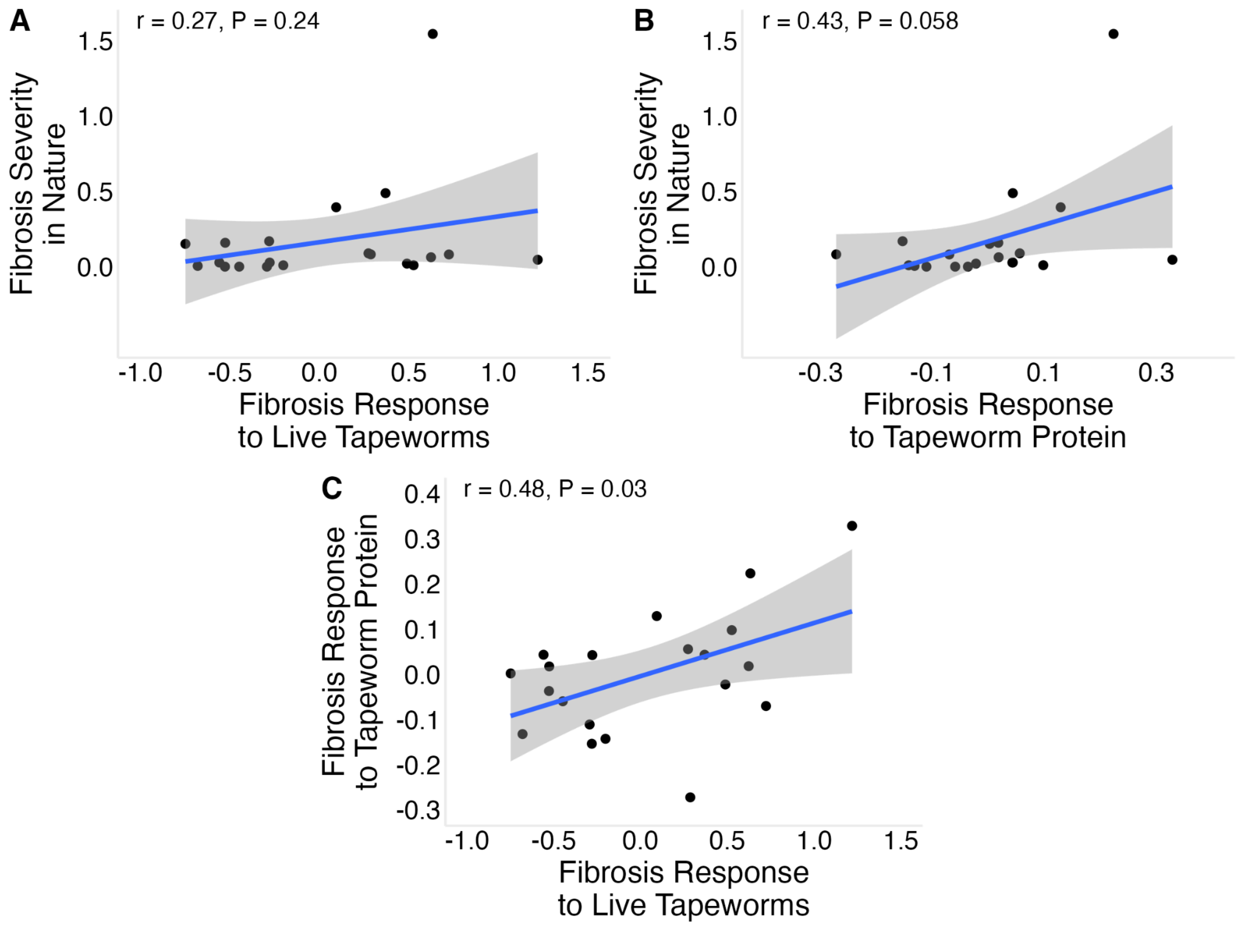


Figure S3: A) Correlation between fibrosis response to live worms and fibrosis severity in nature. B) Correlation between fibrosis response to tapeworm protein and fibrosis severity in nature. C) Correlation between the fibrosis response to live echo tapeworms and the fibrosis response to tapeworm protein. Population-specific fibrosis responses were estimated as posterior mean effect sizes from Bayesian ordinal logistic regression.

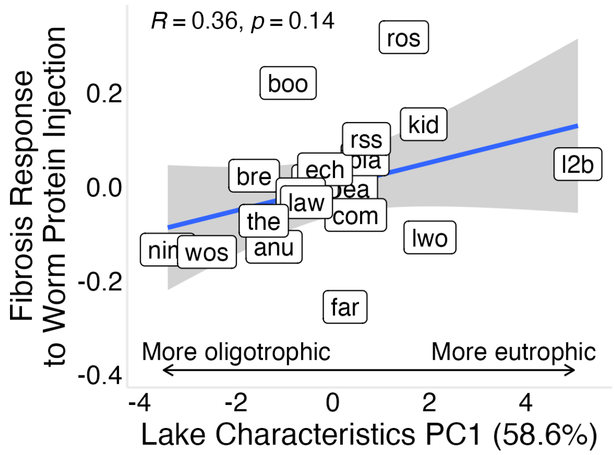


Figure S4: Bayesian estimates of population-level fibrosis response to live tapeworm exposure. Points show posterior mean effect sizes for each population. Effect sizes represent the estimated change in fibrosis score following exposure to live tapeworms relative to controls from a Bayesian ordinal logistic regression model.

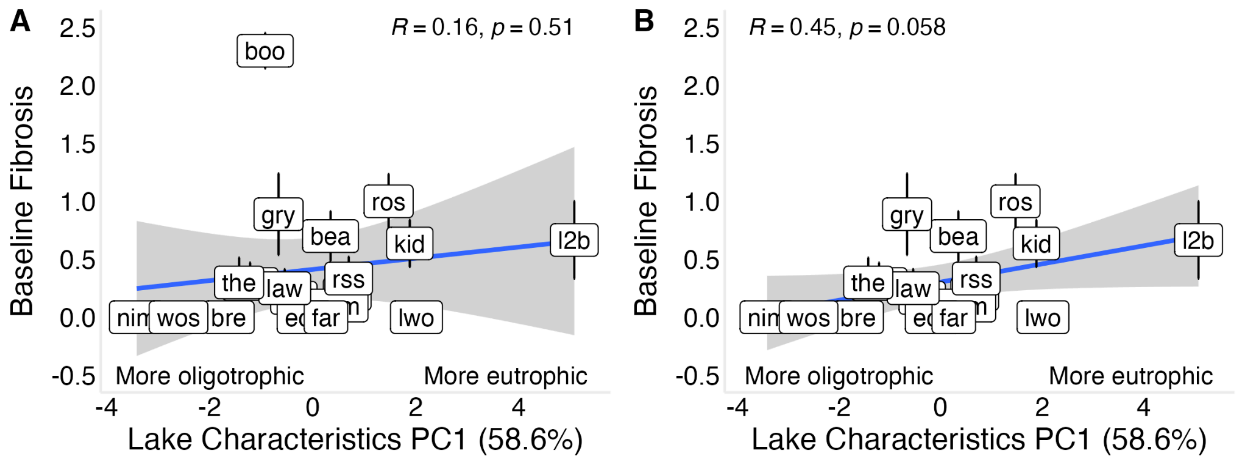


Figure S5: Lake characteristics (PC1) and baseline fibrosis were not significantly correlated when all populations were included (Figure A). However, this relationship became positive and statistically significant after exclusion of Boot Lake (Figure B). PC1 separates more oligotrophic lakes (deeper, higher oxygen, lower chlorophyll *a*) from more eutrophic lakes (shallower, lower oxygen, higher chlorophyll *a*).

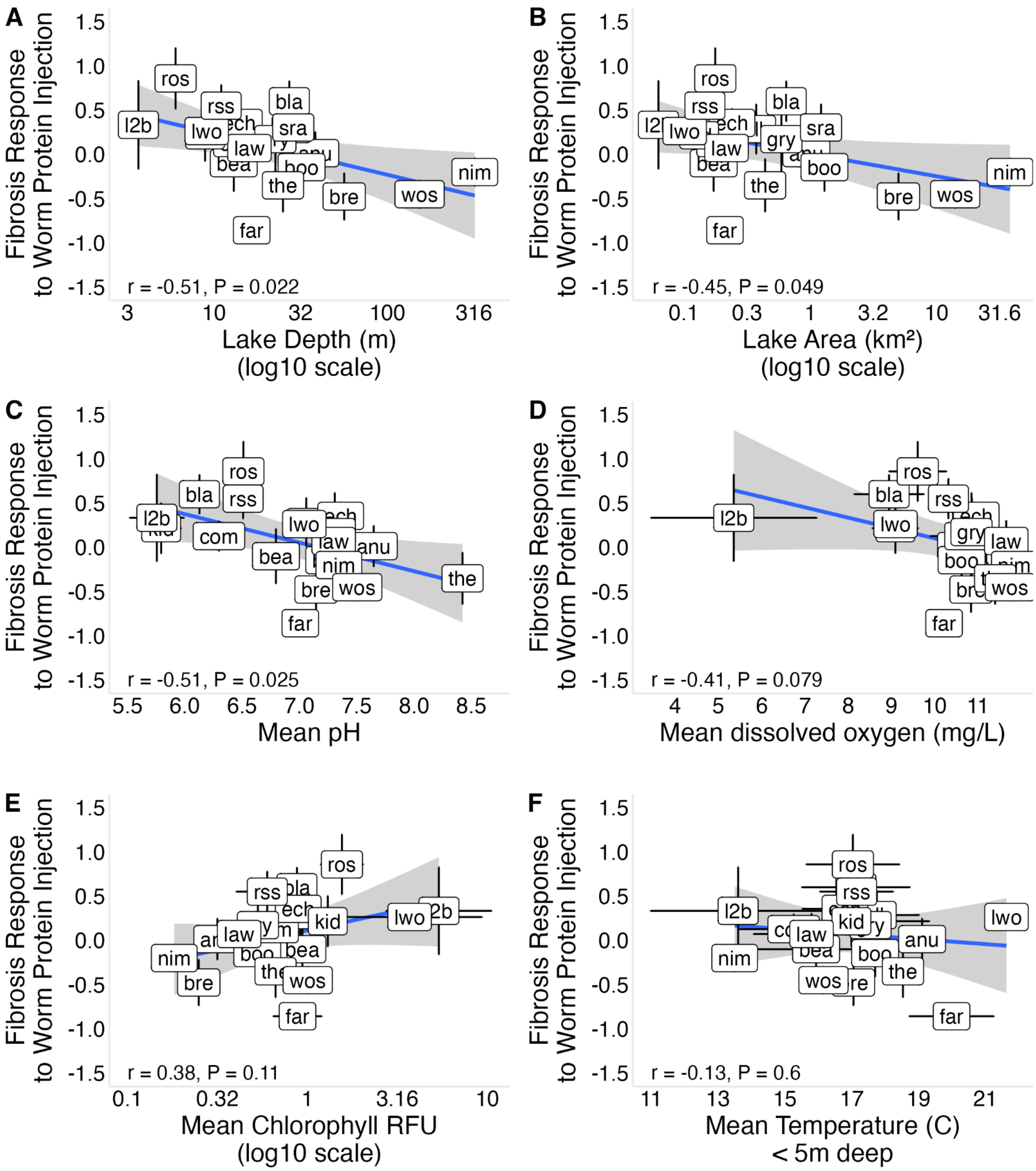


Figure S6: Correlations between ecological covariates and the fibrosis response to tapeworm protein injection, calculated as the difference between fibrosis severity in fish injected with tapeworm protein and fish injected with saline solution. (A) log lake depth (B) log lake area (C)

mean pH (D) mean dissolved oxygen (E) log mean chlorophyll *a* (F) Mean temperature less than 5m deep.

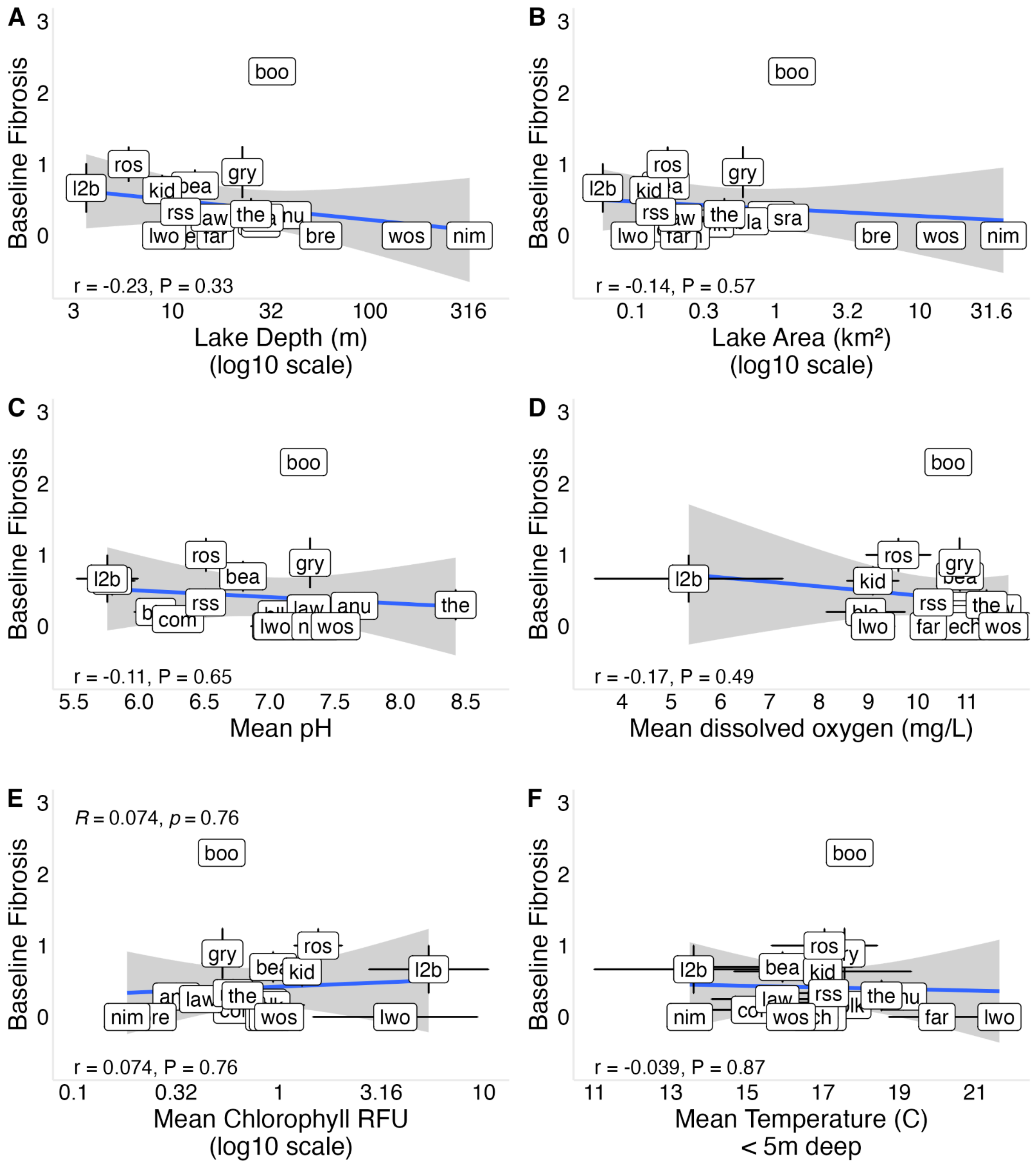


Figure S6: Baseline fibrosis is not correlated with ecological covariates (A) log lake depth (B) log lake area (C) mean pH (D) mean dissolved oxygen (E) log mean chlorophyll *a* (F) mean temperature less than 5m deep.

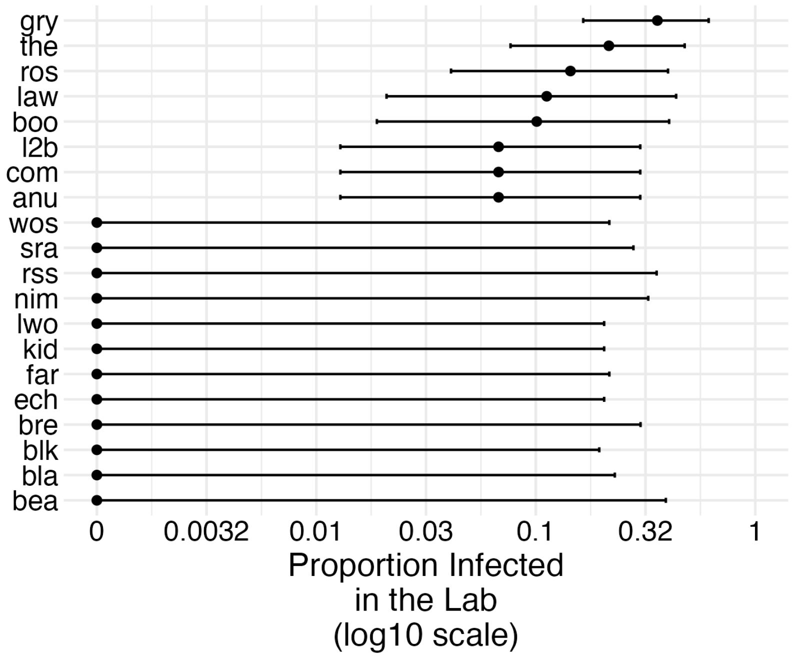


Figure S7: Stickleback populations differ in tapeworm infection success (binomial GLM; population deviance 17.9420, df = 21, P < 0.0001). Plot shows the proportion of exposed fish that were infected for each population along with 95% binomial confidence intervals for the proportion. X-axis is plotted on a log10 scale, however the labels are showing the unlogged values.

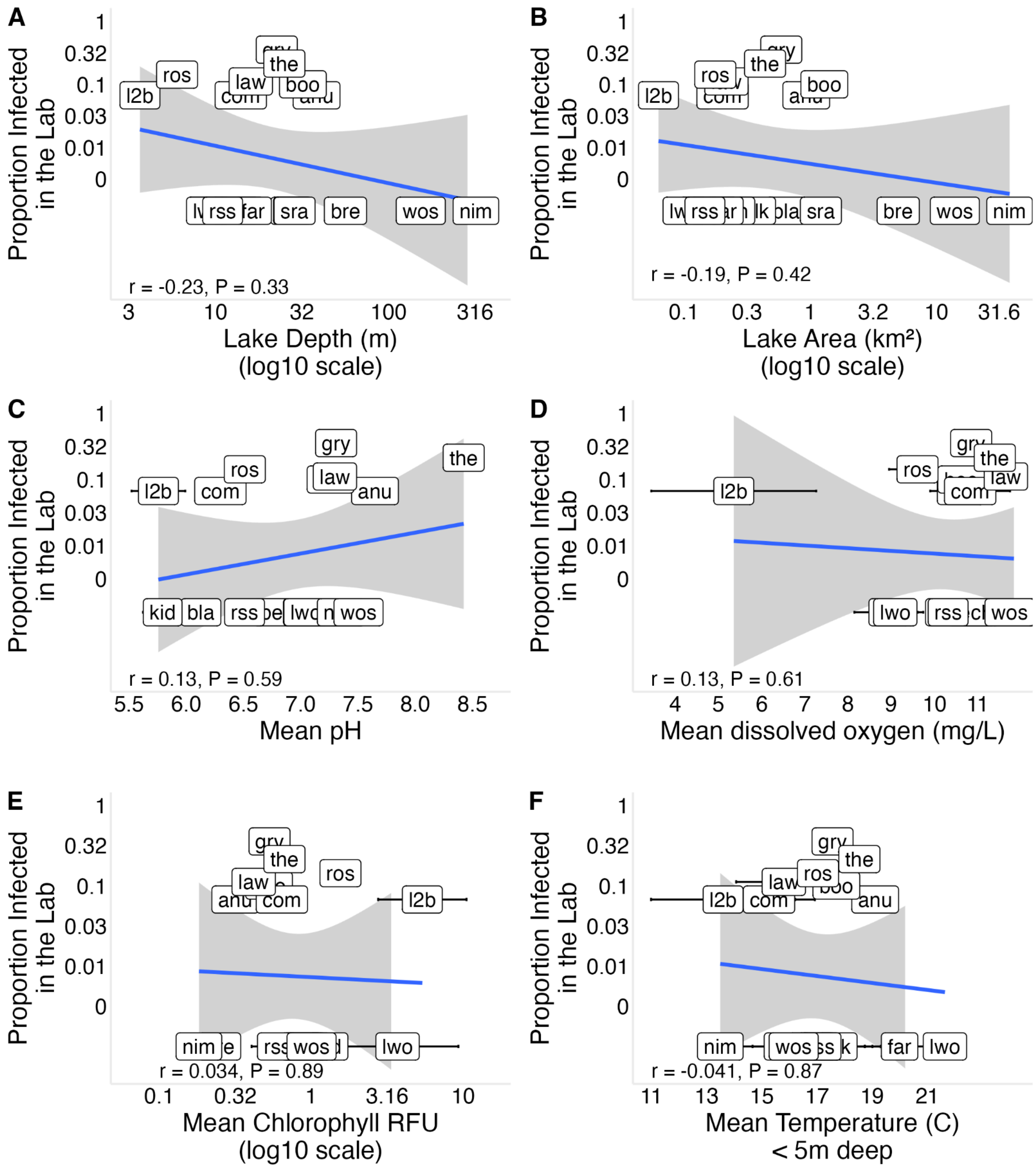


Figure S8: The log proportion of fish infected with tapeworms in the lab is not correlated with ecological covariates (A) log lake depth (B) log lake area (C) mean pH (D) mean dissolved oxygen (E) mean chlorophyll *a* (F) mean temperature less than 5m deep. The Y axes are plotted on a log scale, but labeled with the unlogged values.

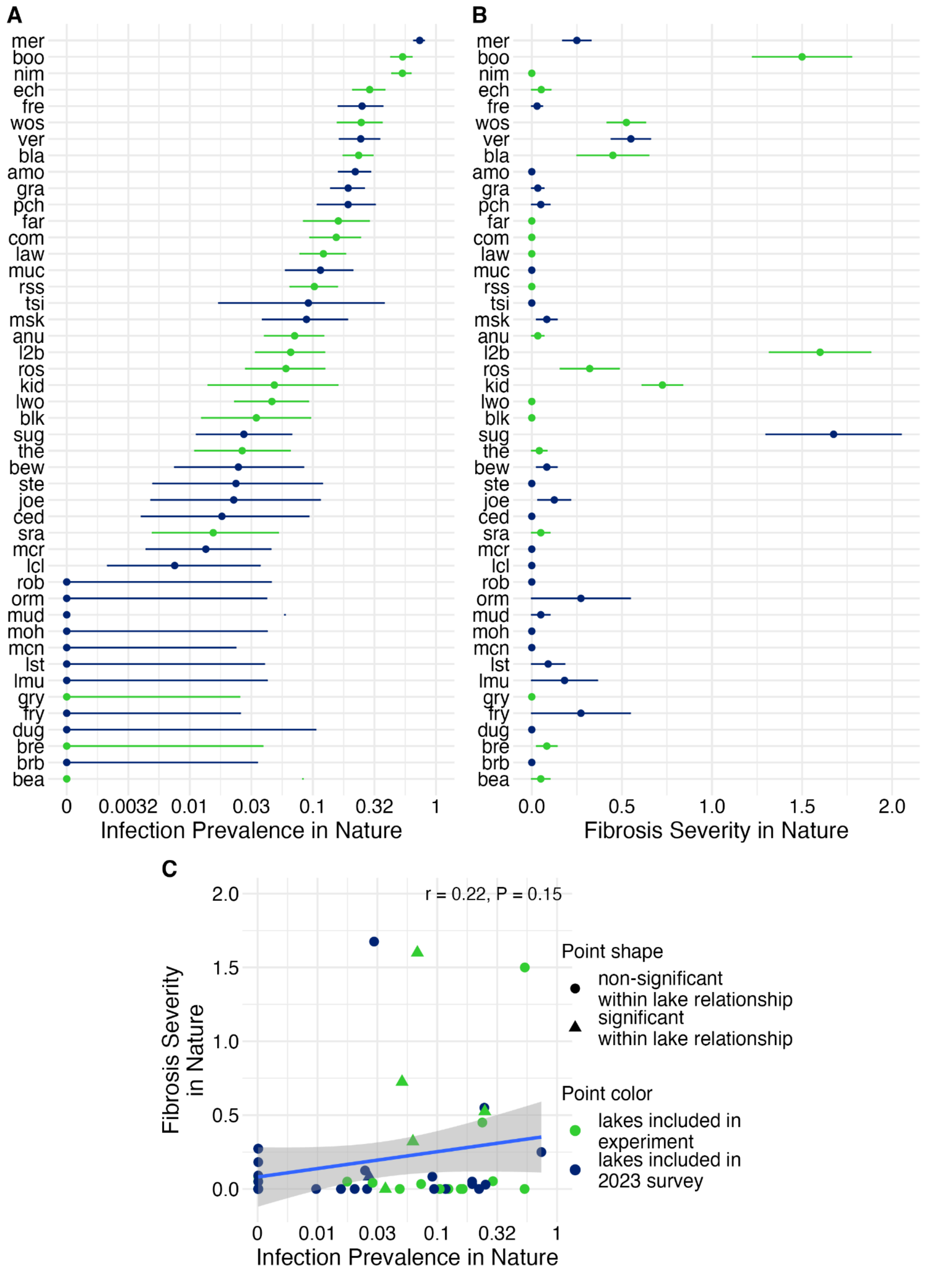


FigS9: Lakes were originally selected with the data represented in this version of Figure 2. Infection prevalence was estimated using approximately 150 fish per lake preserved in ethanol and fibrosis severity was estimated using 10-20 per lake dissected immediately after

euthanasia. Figure 2 was updated using data from approximately 110 fresh and formalin preserved fish per lake.

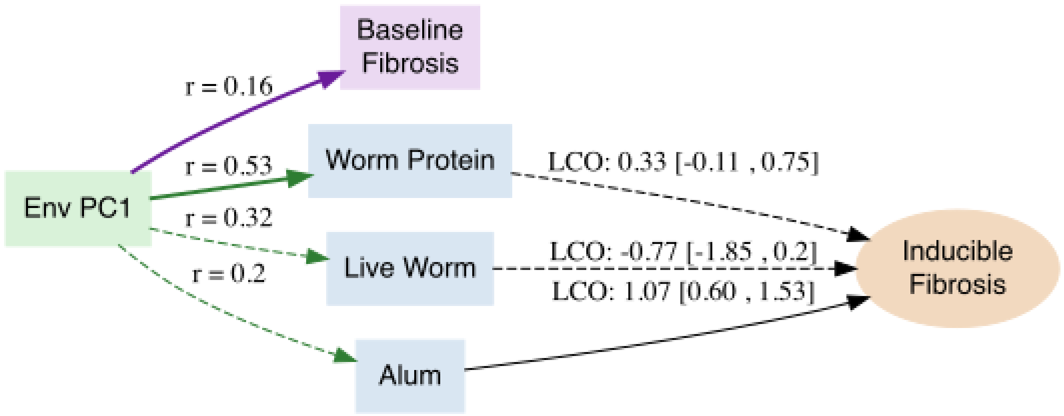


FigS10: Strength of correlation (r) between environmental PC1 and baseline fibrosis, worm protein injection, live worm exposure, or alum injections (bold indicates significant correlation, P<0.05). The Bayesian estimated log cumulative odds for the inducible change in fibrosis with 95% CIs.

| Variable | PC1_loading |
| --- | --- |
| log_max_lake_depth | -0.479 |
| log_area | -0.443 |
| mean_ph | -0.399 |
| mean_o2 | -0.479 |
| mean_temp | 0.005 |
| mean_chlorophyll | 0.431 |

Table S3: Loadings of lake environmental variables on the first principal component (PC1) from a PCA of lake characteristics. PC1 explains 58.6% of the variance.
