## Supplementary_Data2_Bayesian_Posterior_Outputs for "Constitutive and inducible fibrosis explain immune variation among threespine stickleback populations"

### Supplementary Data 2

**This supplement presents the posterior probability distributions from the Bayesian Ordinal Logistic Regressions of fibrosis, as described in the main text. Each page of this supplement corresponds to a different BayesOLR model, showing the posterior distributions of particular interest. Vertical lines represent estimates of zero effect (dashed black line) and the mean estimate (solid blue line). The model and data used for each analysis are described besides each set of distributions.**

A) Strong mong-lake variation in fibrosis in wild caught samples

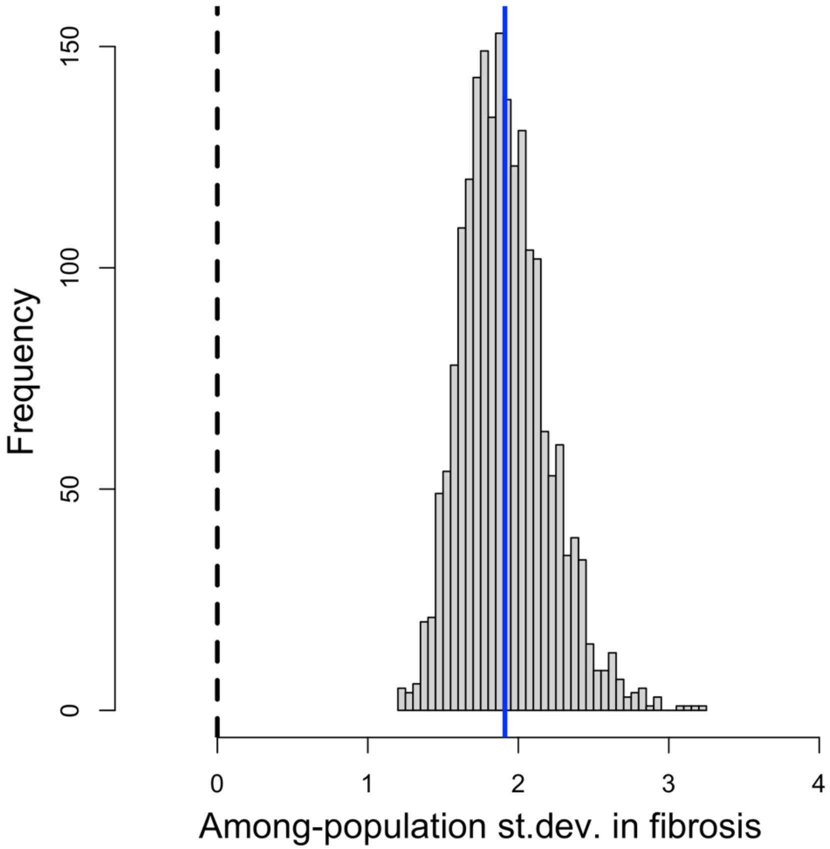


**Bayesian Ordinal Logistic Regression:**

Fibrosis rank ~ Population (random effect, panel A)

**Data**: Lab raised stickleback, unexposed and uninjected

B) Higher fibrosis in infected than uninfected wild caught individuals

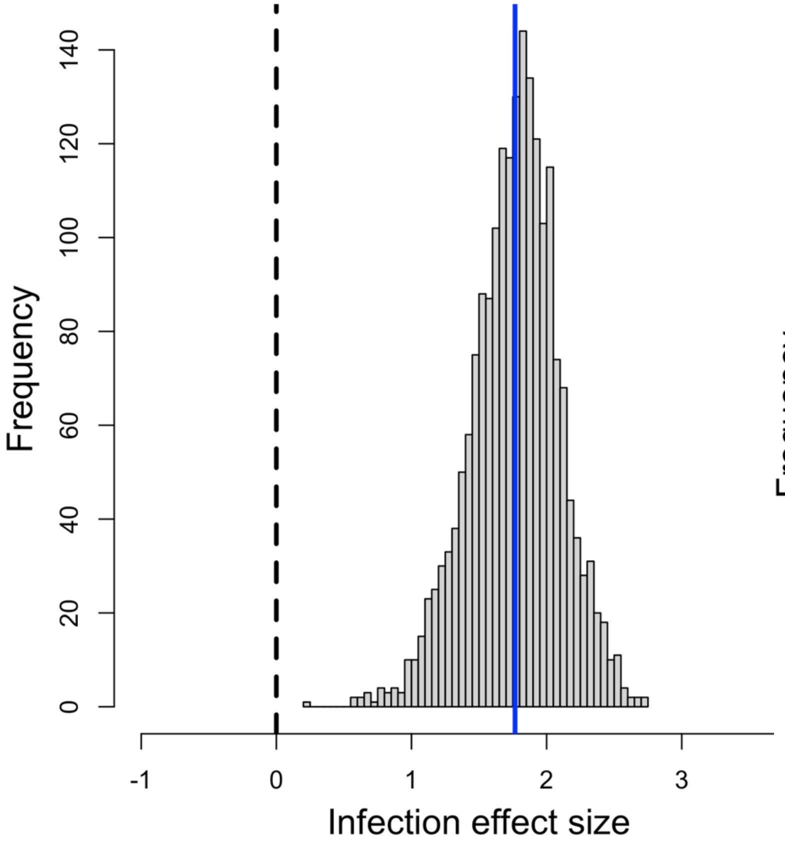


C) Strong among-lake variation in infection-fibrosis relationship

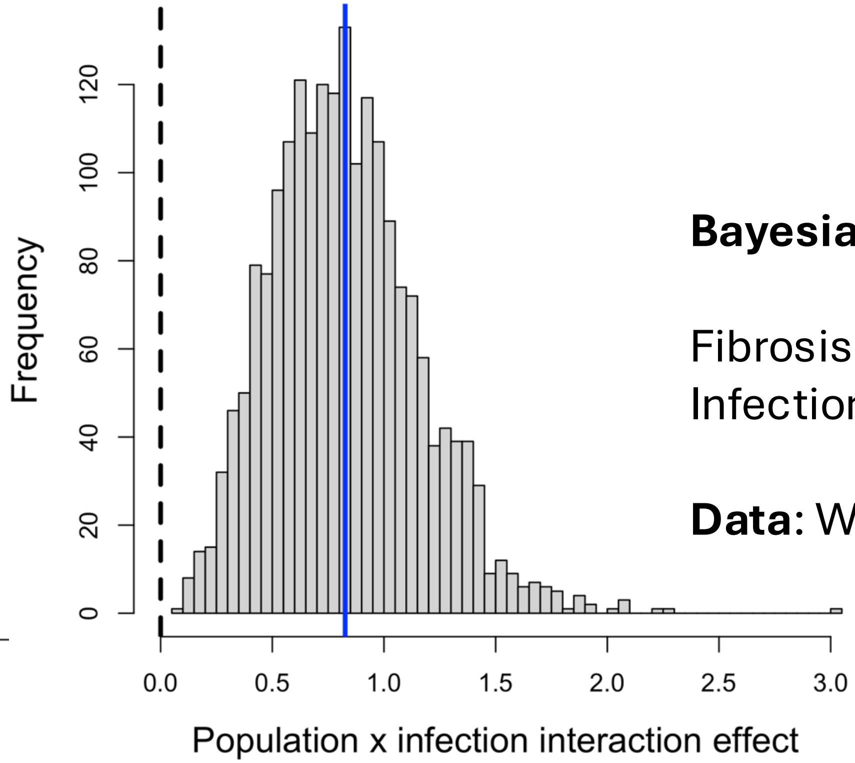


**Bayesian Ordinal Logistic Regression:**

Fibrosis rank ~ Population (random effect) + Infection (B) + Infection*Population interaction (C)

**Data**: Wild-caught stickleback

D) Among-population variation in baseline fibrosis in unchallenged lab fish

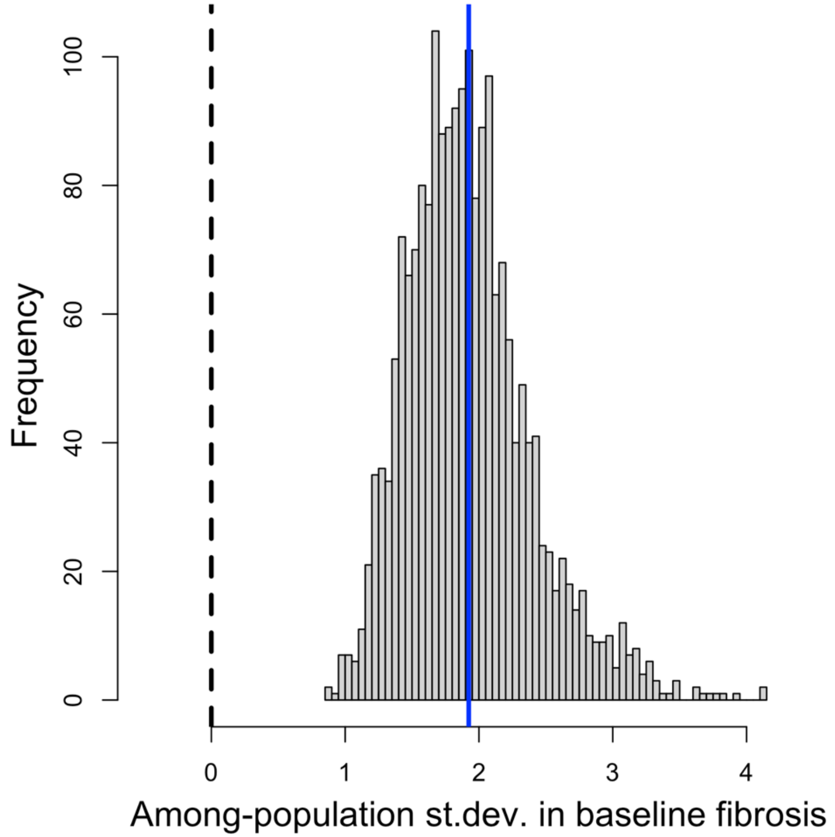


**Bayesian Ordinal Logistic Regression:**

Fibrosis rank ~ Population (random effect, panel D)

**Data**: Lab raised stickleback, unexposed and uninjected

E) Higher fibrosis in Alum-injected than PBS-injected lab fish

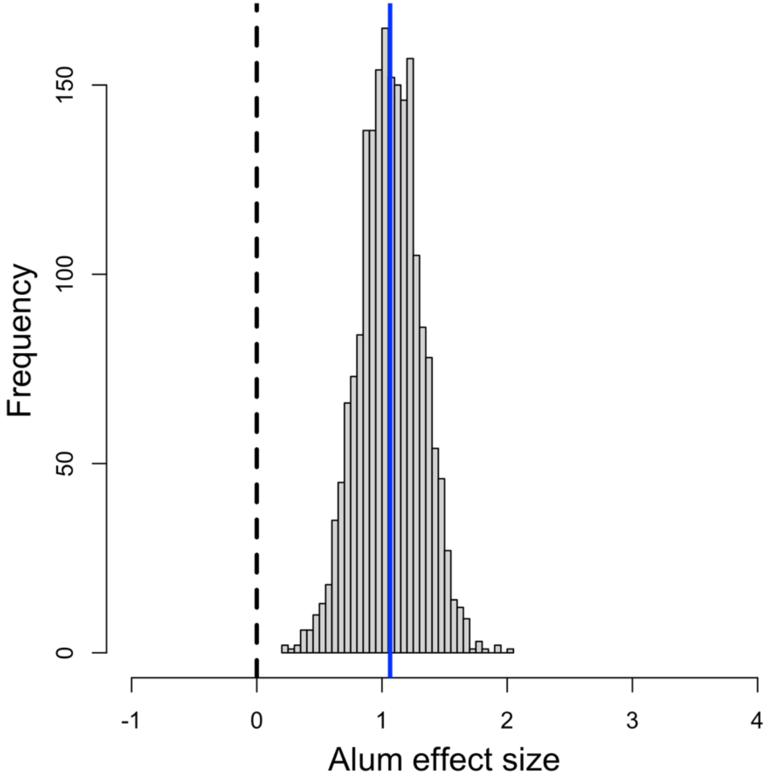


F) Weak among-population variation in injection effect

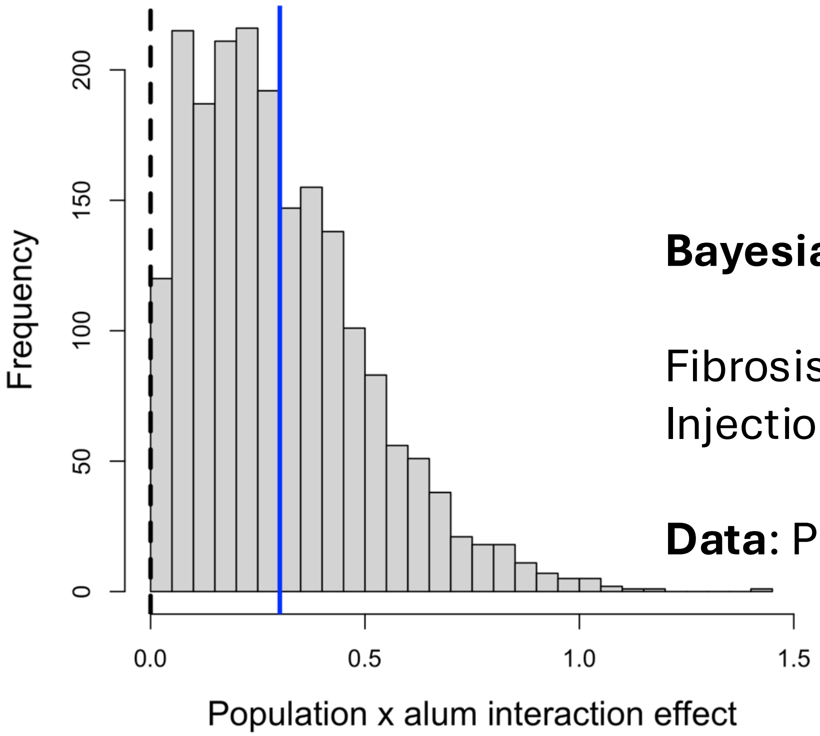


**Bayesian Ordinal Logistic Regression:**

Fibrosis rank ~ Population (random effect) + Injection (E) + Injection*Population interaction (F)

**Data**: PBS-injected and Alum-injected fish (day 2)

G) Higher fibrosis in Alum-injected than PBS-injected lab fish

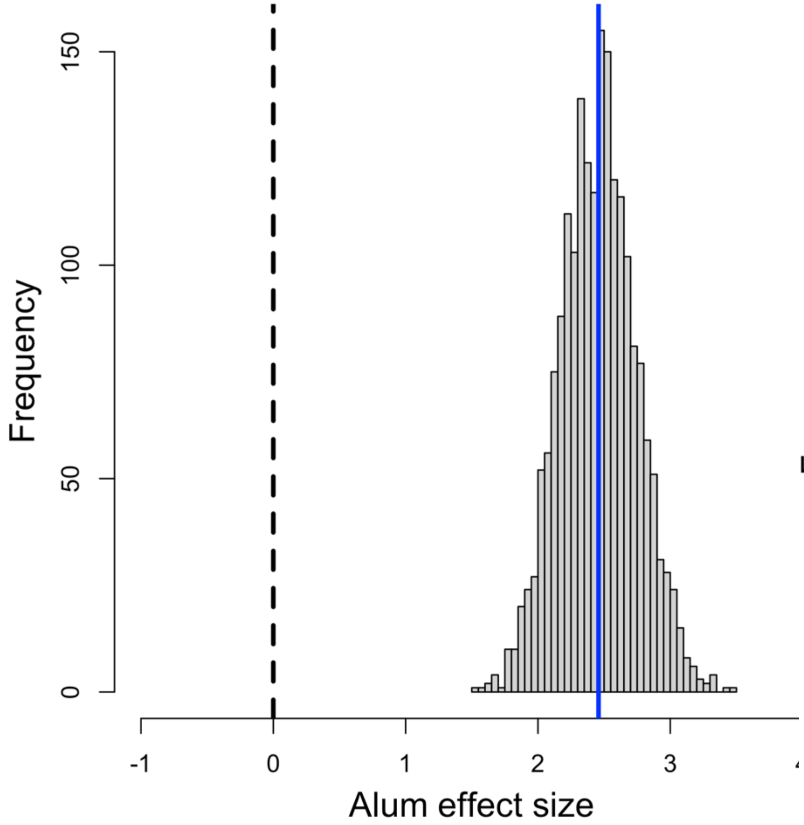


H) Moderate among-population variation in injection effect

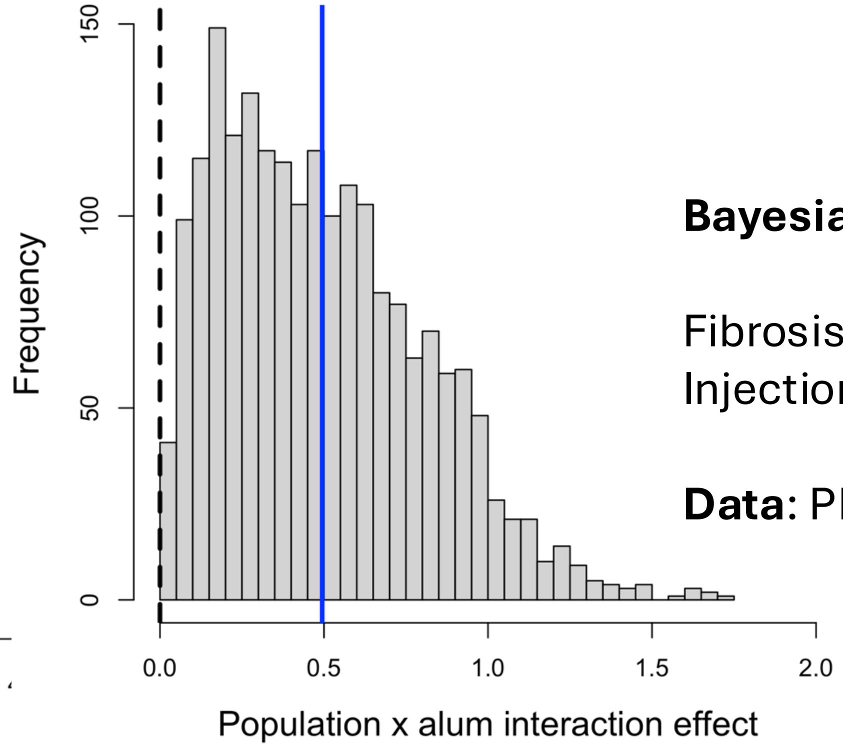


**Bayesian Ordinal Logistic Regression:**

Fibrosis rank ~ Population (random effect) + Injection (G) + Injection*Population interaction (H)

**Data**: PBS-injected and Alum-injected fish (day 7)

I) Fibrosis increases with time in alum-injected lab fish

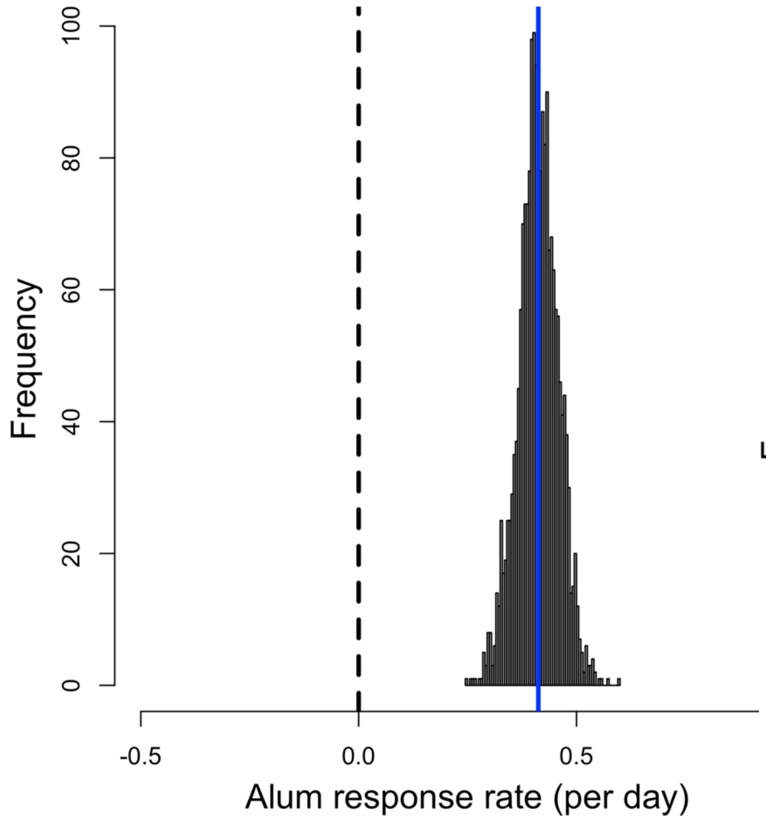


J) Strong among-population variation in rate of fibrosis response to alum

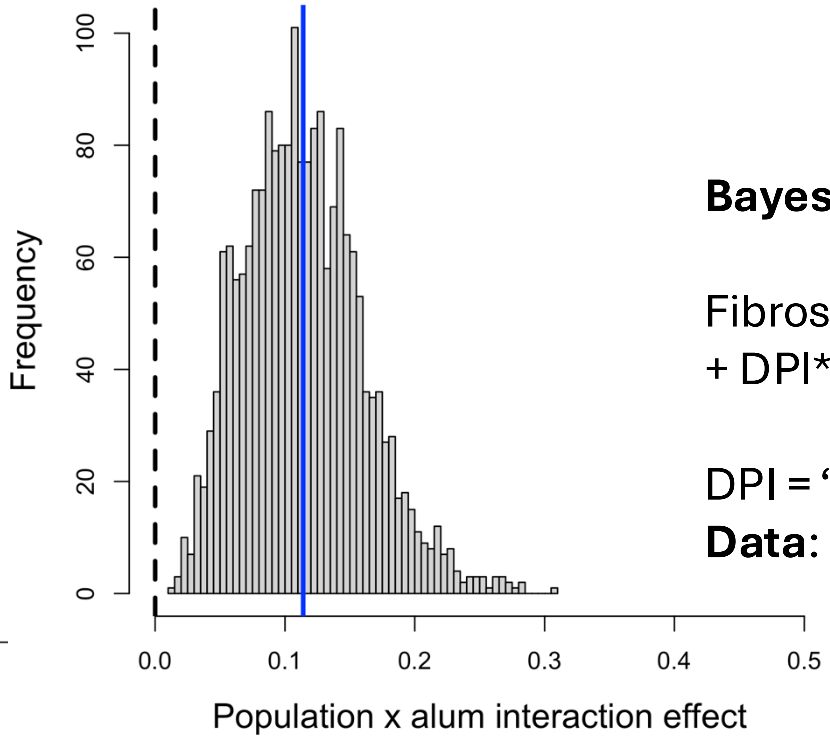


**Bayesian Ordinal Logistic Regression:**

Fibrosis rank ~ Population (random effect) + DPI (I) + DPI*Population interaction (J)

DPI = "Days Post-Injection": 0, 2, or 7
**Data**: PBS-injected and Alum-injected fish

K) Trend towards positive fibrosis response to tapeworm protein inject in lab fish

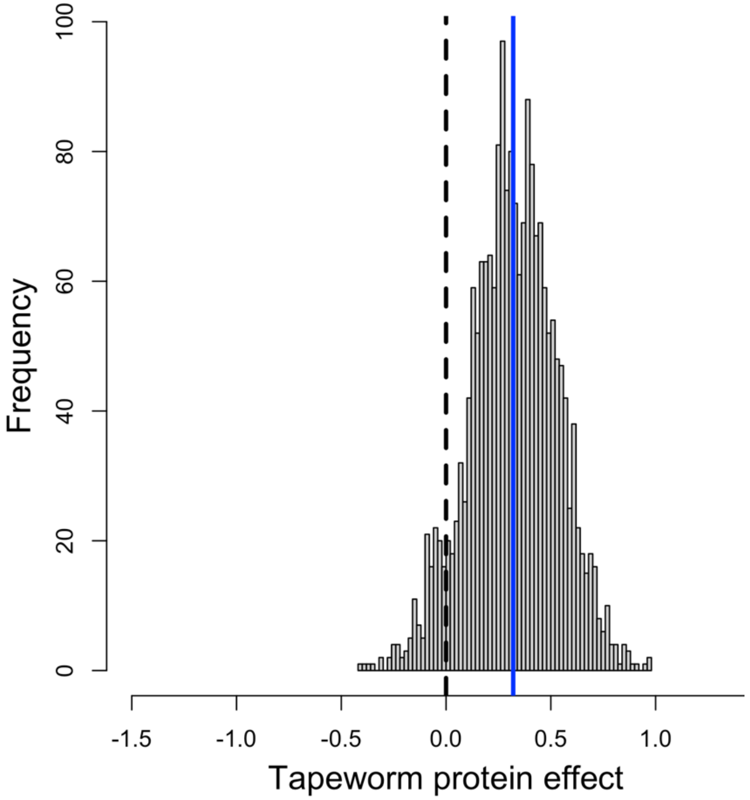


L) Strong support for among-population variation in response to tapeworm protein

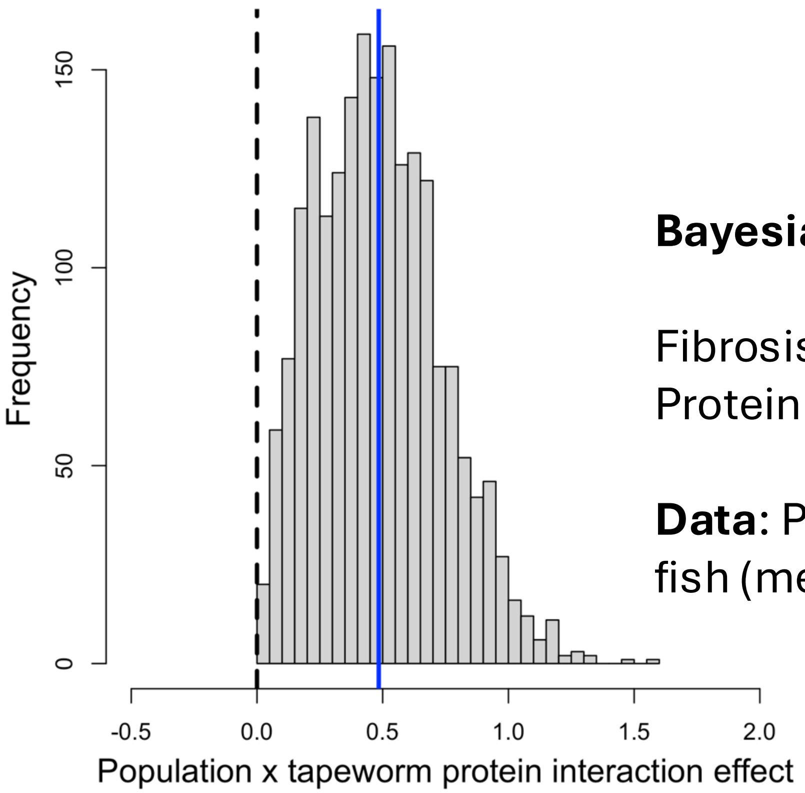


**Bayesian Ordinal Logistic Regression:**

Fibrosis rank ~ Population (random effect) + Protein (K) + Protein*Population interaction (L)

**Data**: PBS-injected and tapeworm protein injected fish (merging local and foreign tapeworms).

M) No overall difference in fibrosis for foreign vs local tapeworm protein in lab fish

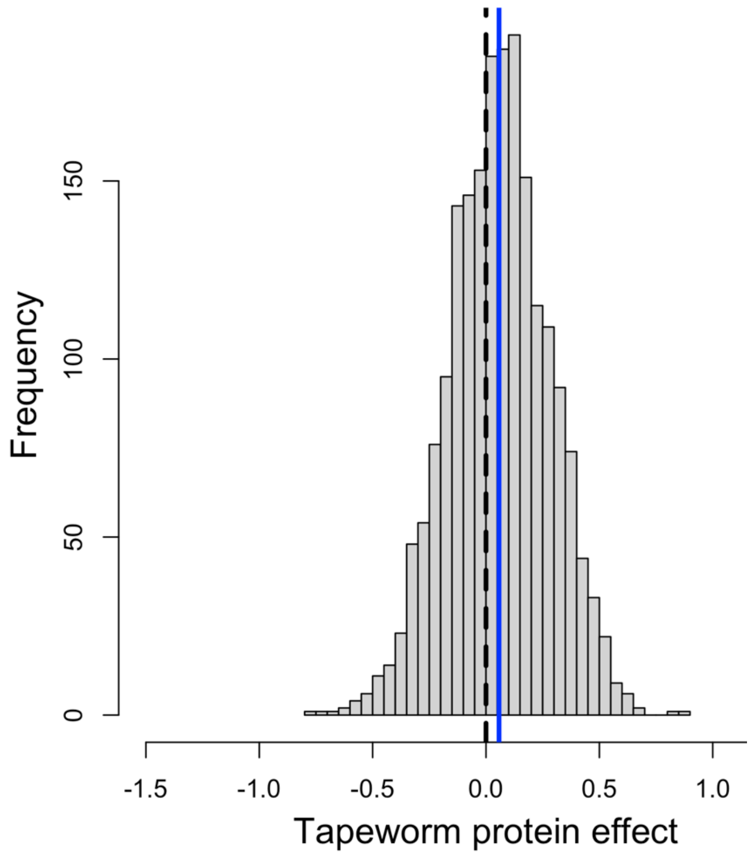


N) Support for weak among-population variation in response to local versus foreign tapeworm protein

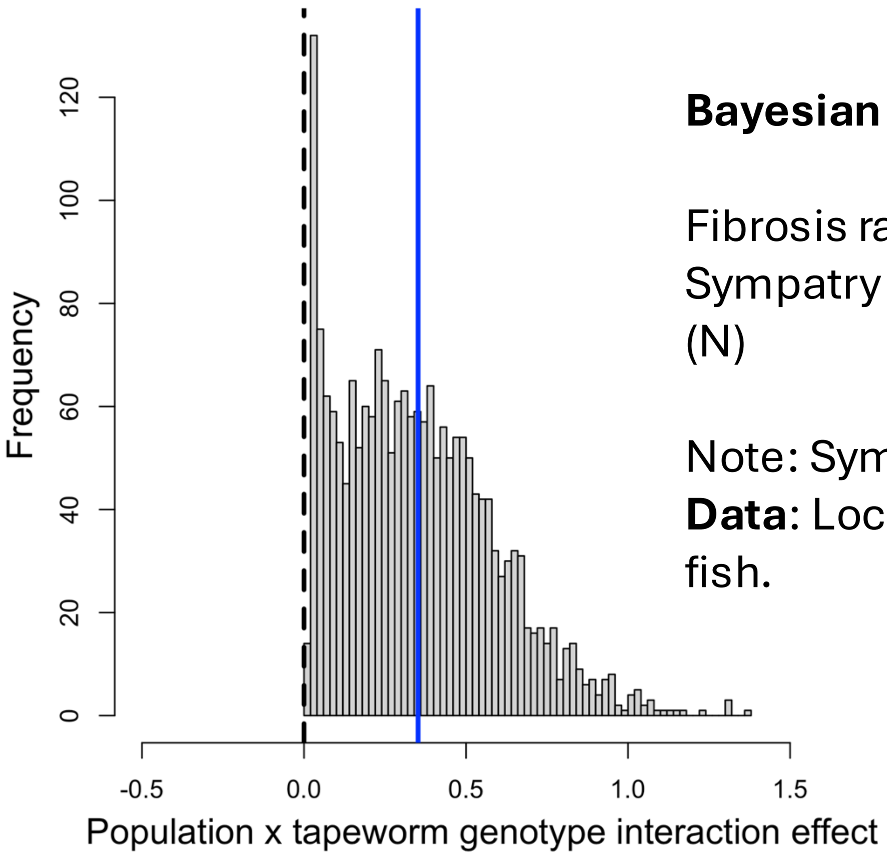


**Bayesian Ordinal Logistic Regression:**

Fibrosis rank ~ Population (random effect) + Sympatry (M) + Sympatry*Population interaction (N)

Note: Sympatry = Local – Foreign protein contrast
**Data**: Local vs foreign tapeworm protein injected fish.

O) Exposure to Echo lake tapeworms induces less fibrosis than hybrid tapeworm exposure

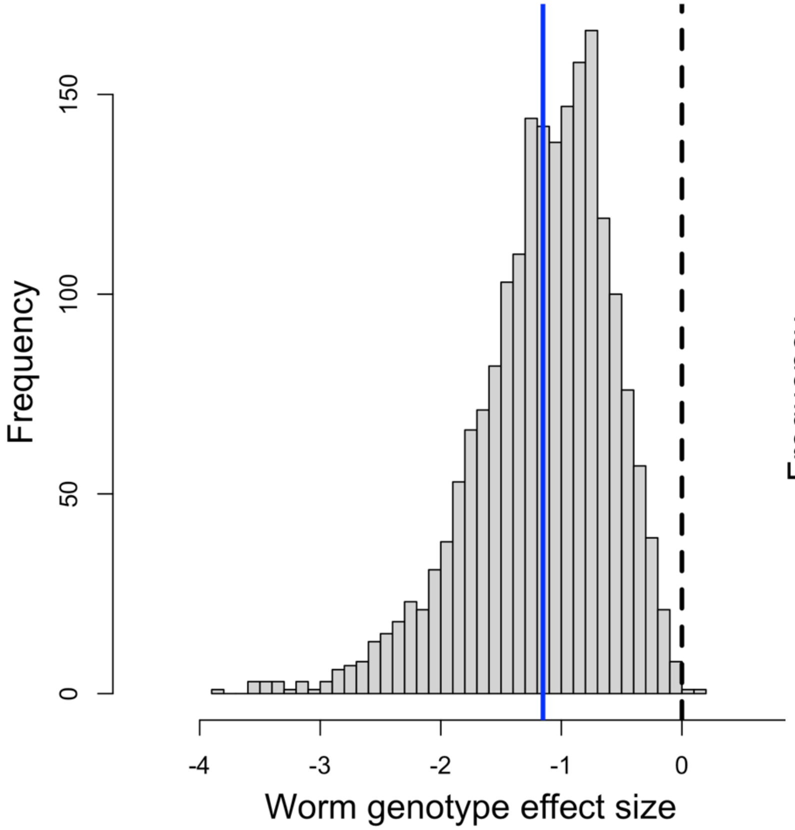


P) Strong support for among-population variation in response to tapeworm genotype

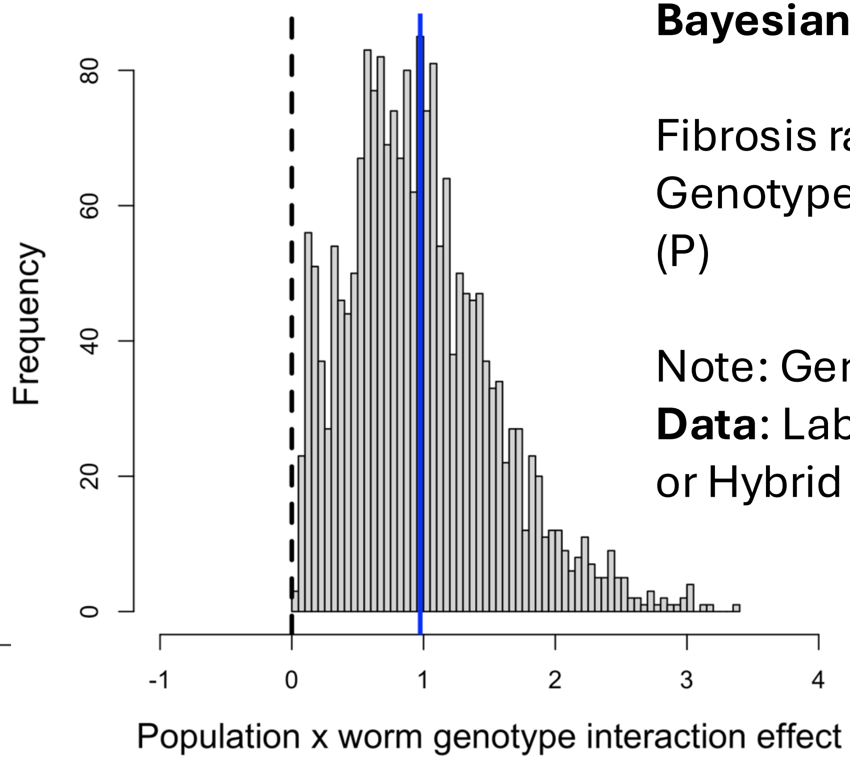


**Bayesian Ordinal Logistic Regression:**

Fibrosis rank ~ Population (random effect) + Genotype (O) + Genotype*Population interaction (P)

Note: Genotype = Echo Lake – Hybrid

**Data**: Lab raised fish exposed to either Echo Lake or Hybrid (Echo x Skogseidvatnet F2s) tapeworms.

Q) Exposure to live Echo Lake tapeworm induces fibrosis

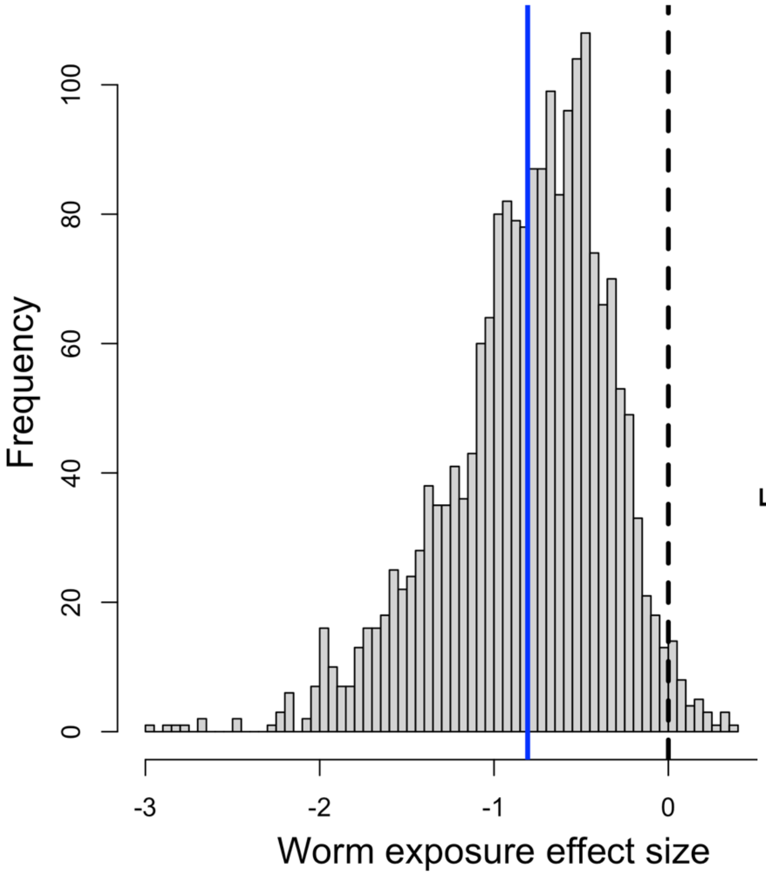


R) Modest support for among-population variation in response to Echo Lake tapeworm exposure

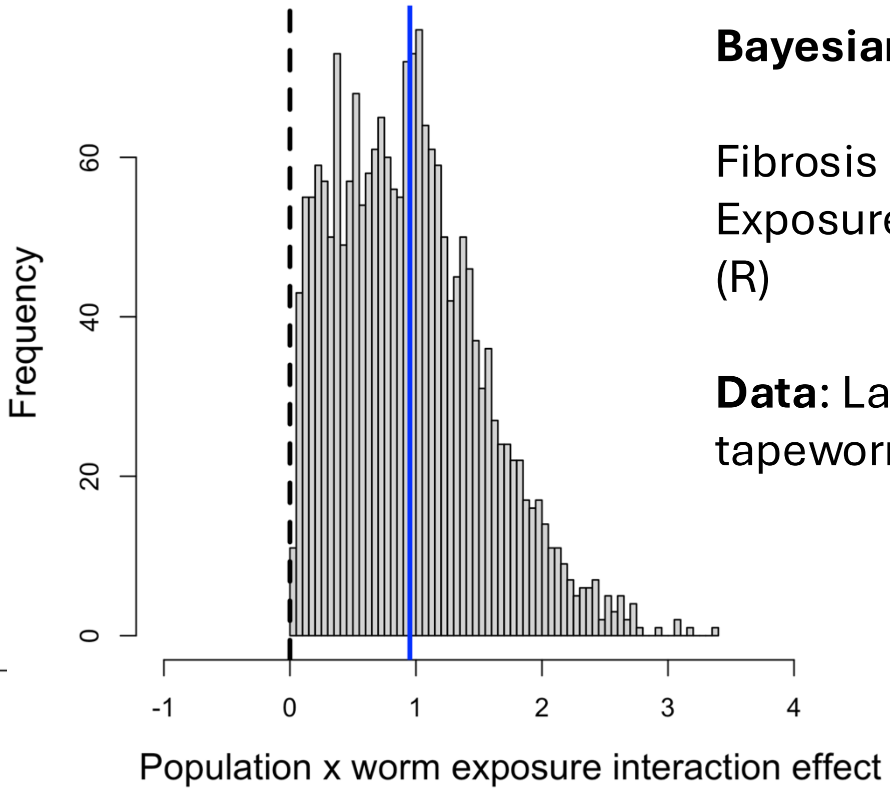


**Bayesian Ordinal Logistic Regression:**

Fibrosis rank ~ Population (random effect) + Exposure (Q) + Exposure*Population interaction (R)

**Data**: Lab raised fish exposed to Echo Lake tapeworms, and unexposed controls.

S) Exposure to live F2 hybrid tapeworms induces no consistent fibrosis response

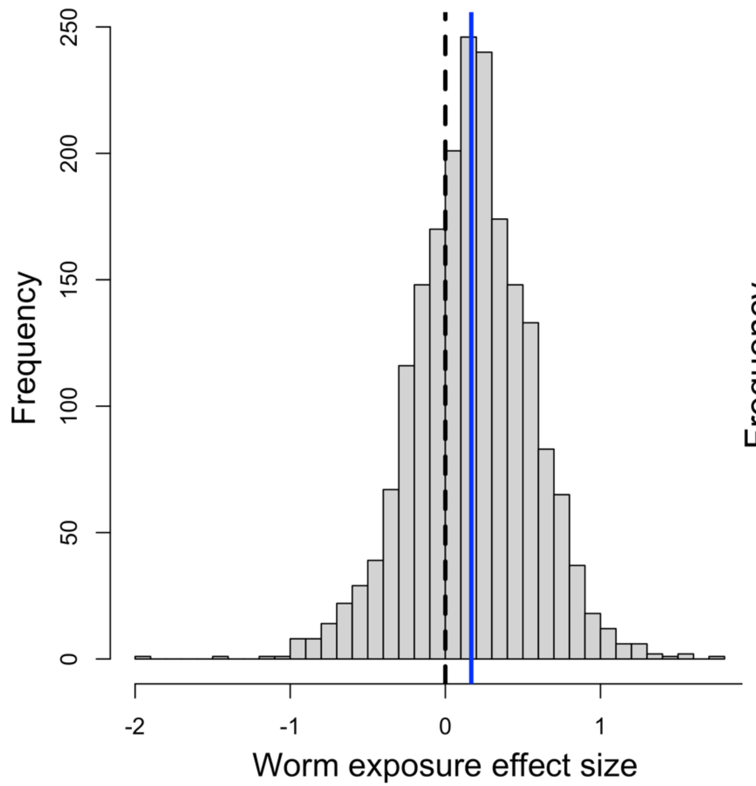


T) Strong support for among-population variation in response to F2 hybrid tapeworm exposure

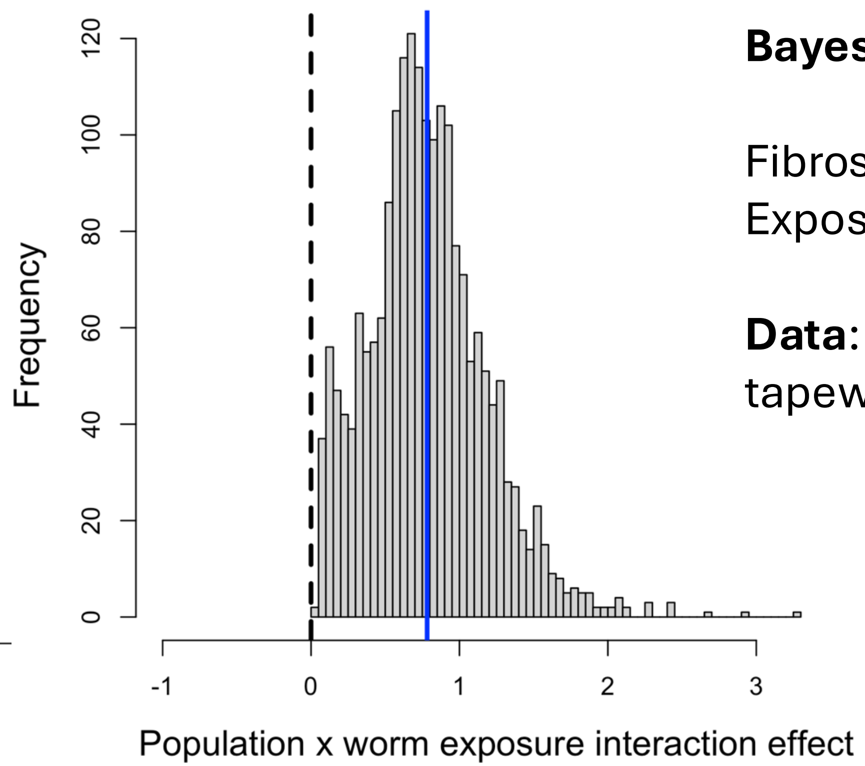


**Bayesian Ordinal Logistic Regression:**

Fibrosis rank ~ Population (random effect) + Exposure (S) + Exposure*Population interaction (T)

**Data**: Lab raised fish exposed to F2 hybrid tapeworms, and unexposed controls.

S) Exposure to live F2 hybrid tapeworms induces no consistent fibrosis response

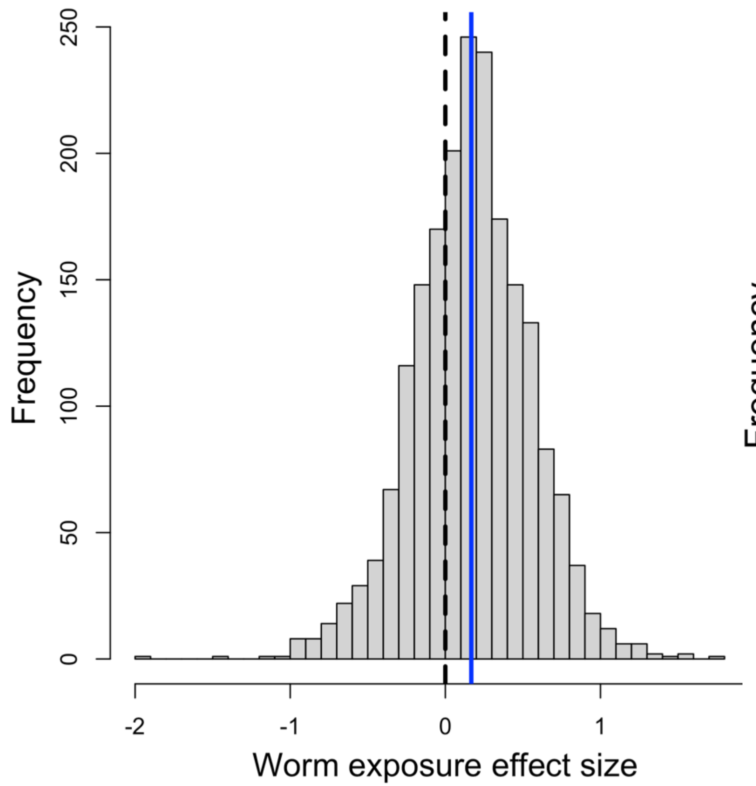


T) Strong support for among-population variation in response to F2 hybrid tapeworm exposure

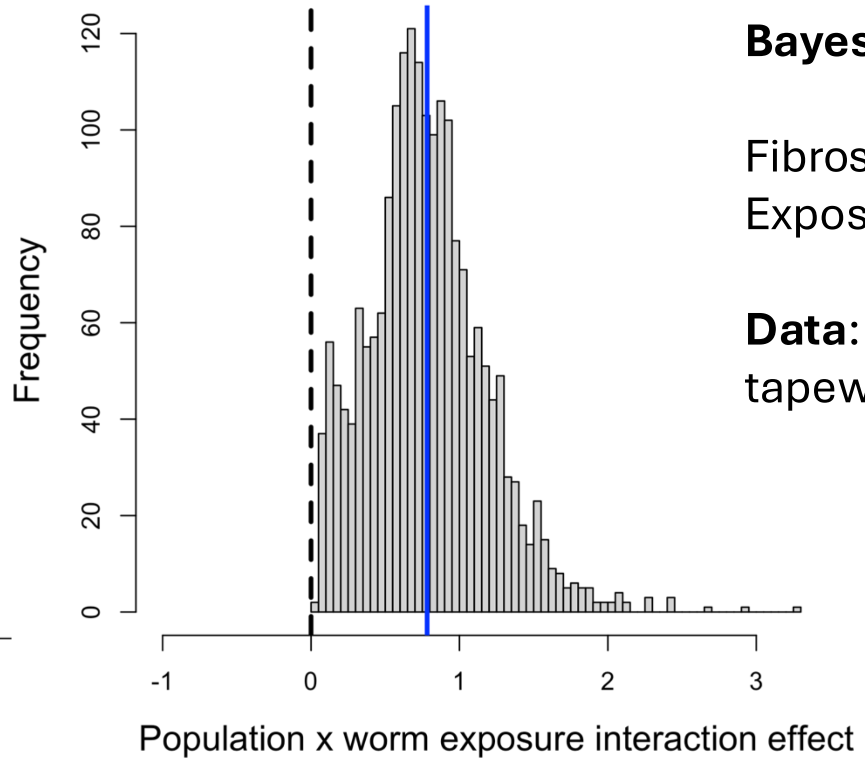


**Bayesian Ordinal Logistic Regression:**

Fibrosis rank ~ Population (random effect) + Exposure (S) + Exposure*Population interaction (T)

**Data**: Lab raised fish exposed to F2 hybrid tapeworms, and unexposed controls.
